# NudC moonlights in ribosome biogenesis and homeostasis in *Drosophila melanogaster* polyploid cells

**DOI:** 10.1101/2025.09.14.675990

**Authors:** Duoduo Shi, Yuko Shimada-Niwa, Naoki Okamoto, Akira Nakamura, Yuya Ohhara, Wei Sun, Ryusuke Niwa

## Abstract

Ribosomes, the cellular machinery responsible for protein synthesis, are fundamental across all kingdoms of life. Disruption in ribosome biogenesis (RiBi) can cause severe ribosomopathies, underscoring the critical need for precise regulatory mechanisms governing RiBi. In this study, we identified the gene *NudC* (*nuclear distribution C, dynein complex regulator*), a previously unrecognized regulator of RiBi in polyploid cells of *Drosophila melanogaster* larvae. RNAi-mediated depletion of *NudC* in polyploid salivary gland cells led to a significant reduction in ribosome abundance, accompanied by the loss of ribosome-binding sites on rough endoplasmic reticulum and impaired translation. These defects are linked to decreased levels of nucleolar ribosomal RNA. Notably, *NudC* knockdown also triggered a homeostatic response, characterized by increased transcription and translation of both ribosome biogenesis factors (RBFs) and ribosomal proteins. This response parallels that seen in RBF-deficient cells, suggesting that NudC and RBFs cooperate to maintain RiBi homeostasis. Meanwhile, *NudC*-deficient cells exhibited chromosome abnormalities, activated JNK signaling, and underwent autophagy, closely resembling the defects observed upon loss of RBFs. Finally, our findings suggest that the role of NudC in RiBi is independent of its established function in dynein regulation, indicating the moonlighting role in RiBi played by this gene. Together, these results uncover a new, fundamental function for NudC in maintaining RiBi and homeostasis in polyploid cells, with broader implications for understanding conserved mechanisms of NudC function and RiBi across species.

## Introduction

Ribosomes are essential ribonucleoprotein complexes that translate messenger RNA (mRNA) into proteins, a process fundamental to cell growth and metabolism across all forms of life (Kang et al., 2021). The assembly of ribosomes, known as ribosome biogenesis (RiBi), is a central process that ensures proper protein synthesis and overall cellular function.

In eukaryotes, RiBi occurs primarily in the nucleolus. Ribosomal DNA (rDNA) is transcribed by RNA polymerase I into a 47S precursor ribosomal RNA (rRNA), which contains the sequences of 18S, 5.8S, and 28S rRNAs (Lafontaine, 2015; Pelletier et al., 2018). This precursor is incorporated into the 90S processome, alongside the 5S rRNA and ribosomal proteins (RPs). The 5S rRNA is transcribed in the nucleoplasm by RNA polymerase III, whereas RP mRNAs are synthesized by RNA polymerase II, exported to the cytoplasm for translation, and subsequently imported into the nucleolus. Within the nucleolus, rRNAs and RPs assemble into small (pre-40S) and large (pre-60S) subunits. RiBi is a complex process that depends on numerous ribosome biogenesis factors (RBFs), which coordinate rRNA processing along with the production, modification, and assembly of ribosomal components (Greber, 2016). These steps ultimately result in the production of mature 80S ribosomes capable of sustaining protein synthesis (Pelletier et al., 2018).

Fine-tuned RiBi is essential for maintaining cellular and organismal homeostasis, and disruptions in this process are broadly linked to disease. In humans, defects in RiBi cause a class of disorders collectively termed ribosomopathies, whereas elevated RiBi drives tumor growth (Buszczak, 2023; Khan et al., 2024; Kirby et al., 2015; Nait Slimane et al., 2020; Narla and Ebert, 2010; Zhang et al., 2002). Mechanisms that respond to defects in RiBi have been characterized in various biological systems, particularly those that maintain a balance in transcription, processing, and maturation of rRNA and RPs (Buszczak, 2023; Warner and McIntosh, 2009). In the yeast *Saccharomyces cerevisiae*, feedback mechanisms maintain appropriate RP levels by reducing their synthesis when they accumulate in excess (Eng and Warner, 1991; Gabunilas and Chanfreau, 2016; Roy et al., 2020). Surplus RPs are further eliminated through rapid degradation by the ubiquitin–proteasome system (Sung et al., 2016). Similar regulatory mechanisms have been described in the nematode *Caenorhabditis elegans* and the fruit fly *Drosophila melanogaster*, where some RPs are under feedback control (Noma et al., 2017; Takei et al., 2016; Kitamura et al., 2024). In vertebrates, analogous responses have been reported: in zebrafish, impaired precursor rRNA (pre-rRNA) processing reduces rRNA levels, increasing the expression of RiBi–related genes (Zhu et al., 2021); In aged mouse skeletal muscle, reduced pre-rRNA expression coincides with upregulation of RP transcripts (Kirby et al., 2015).

The fruit fly *D. melanogaster* provides a powerful model for investigating RiBi, owing to its robust genetic tools and the conservation of key cellular processes with other eukaryotes. Mutations in RBFs or RPs in *D. melanogaster* often result in lethality, growth defects, and developmental delays (Brehme, 1939; He et al., 2015). At the tissue level, defects in RiBi can trigger a nucleolar stress response, manifesting as cell cycle arrest, apoptosis, or autophagy (DeLeo et al., 2021). Importantly, different tissues exhibit distinct responses to impaired RiBi. For example, diploid cells in *D. melanogaster*, such as imaginal disc cells, are highly sensitive to disruptions in RiBi, frequently responding with apoptosis (James et al., 2013; Wang and DiMario, 2017). In contrast, polyploid cells, including those in the salivary gland (SG) and midgut, utilize compensatory strategies such as autophagy and metabolic reprogramming to preserve cellular homeostasis. (He et al., 2015; James et al., 2013; Wang and DiMario, 2017). These differential responses illustrate how the functional roles and biosynthetic demands of tissues shape their sensitivity to disruptions in RiBi. Such tissue-specific outcomes suggest that cells deploy distinct regulatory systems to cope with impaired ribosome production. However, the molecular mechanisms that orchestrate these systems—spanning transcriptional, post-transcriptional, and protein-level regulation—remain poorly understood.

In this study, we identified the gene *nuclear distribution C, dynein complex regulator* (*NudC*) as a key regulator of RiBi homeostasis in the polyploid SG cells of *D. melanogaster* larvae. *NudC* is highly conserved across a broad range of species, from fungi to mammals (Riera and Lazo, 2009). Previously, NudC functions as a microtubule-associated protein that regulates cytoplasmic dynein activity, contributing critically to mitotic spindle formation, cytokinesis, nuclear migration, and intracellular transport (Aumais et al., 2001; Islam et al., 2020; Morris et al., 1998; Zhou et al., 2003). However, evidence has suggested that NudC also performs roles independent of dynein trafficking (Garner et al., 2024; Riera et al., 2007), indicating that its full range of functions is yet to be clarified. Here, we reveal a previously unrecognized, dynein-independent role for NudC in maintaining RiBi. Knockdown of *NudC* in SG cells causes reduced rRNA levels and activates compensatory homeostatic pathways that increase the transcription and translation of RBFs and RPs. Notably, the molecular responses in *NudC*-deficient cells closely resemble those observed in cells lacking RBFs, suggesting that NudC collaborates within the RiBi maintenance network. We propose that NudC performs a unique moonlighting function critical for RiBi homeostasis in polyploid cells.

## Results

### NudC maintains cell growth and tissue function in the prothoracic gland

*NudC* was selected for analysis because of its potential role in ecdysteroid biosynthesis, a critical process that regulates molting and metamorphosis in insect development. Ecdysteroids are produced in the prothoracic gland (PG), a specialized endocrine organ, in a tightly regulated spatiotemporal manner (Niwa and Niwa, 2025, 2014). Previous genome-wide RNAi screens in the PGs of *D. melanogaster* identified 701 candidate genes that potentially regulate ecdysteroid biosynthesis based on developmental defects (Danielsen et al., 2016). A subsequent study further narrowed this from 701 to 449 candidate genes (Ohhara et al., 2019). Among these candidates, *NudC* stood out as the sole gene classified as a “microtubule binding motor protein” by the PANTHER classification system (https://pantherdb.org/). While microtubules have been implicated in regulating ecdysone biosynthesis in PGs (Watson et al., 1996), the broader relationship between microtubule-associated proteins and ecdysteroid production remains poorly explored.

We confirmed that knocking down *NudC* in PG cells causes developmental arrest during the third instar larval (L3) stage (Figures 1A and 1B). Three major phenotypes were observed in *NudC*-deficient PG cells. First, the PG size was significantly reduced compared to the controls (Figures 1C and 1D). Secondly, microtubules, visualized by EGFP-tagged β-Tubulin (β-Tubulin::EGFP), accumulated abnormally around the nucleus (Figure S1A). Third, the polytene chromosome structure was altered, with nuclei exhibiting five prominent DNA-enriched “blobs” (Figure 1E). These blobs likely correspond to the five major *D. melanogaster* chromosome arms (X, 2L, 2R, 3L, 3R), a configuration previously reported in developing nurse cells undergoing intermediate endoreplication (Dej and Spradling, 1999; Volpe et al., 2001; Zhimulev and Koryakov, 2009). Endoreplication allows cells to increase in size by repeatedly replicating their DNA without cell division, resulting in polyploid cells. Our findings suggest that *NudC* knockdown disrupts the normal endoreplication process in PG cells, as evidenced by reduced ploidy levels measured by C-value (the constant value of haploid DNA content) in *NudC*-deficient PGs compared to controls (Figure 1F).

**Figure 1.**
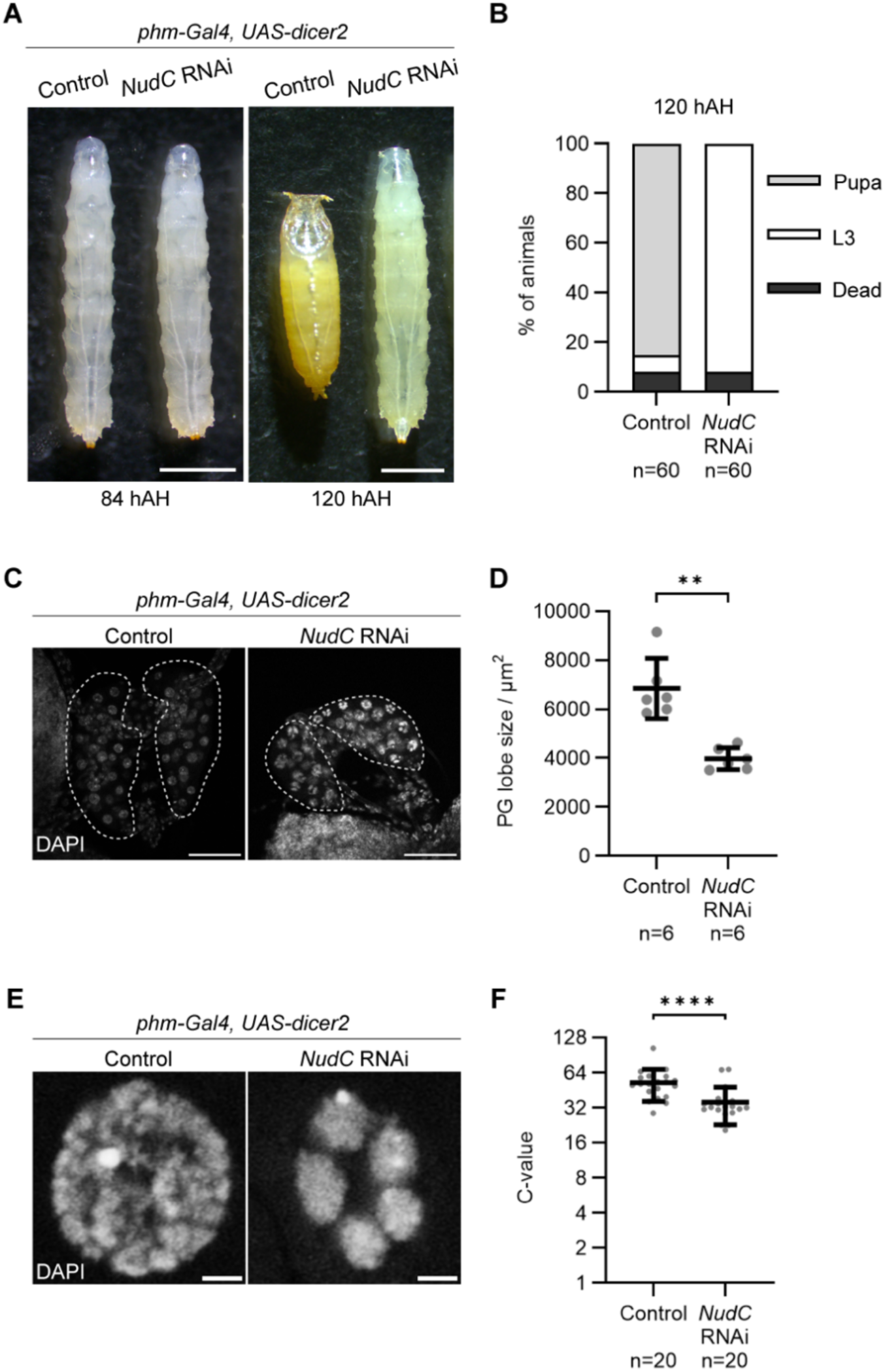
NudC supports tissue growth and maintains chromosomal integrity in prothoracic gland (PG) cells. (A) Representative images of *Drosophila melanogaster* at 84 hAH (left) and 120 hAH (right). Control (*phm*>*dicer2, +*) and *NudC* knockdown (*phm*>*dicer2*, *NudC RNAi*) flies are shown. Scale bar: 1 mm. (B) Survival and developmental progression of control and *NudC* RNAi flies at 120 hAH (n=60). L3 indicates the third-instar larvae. Sample sizes (the number of flies) are shown below each bar. (C) PGs from wandering L3 larvae (89 hAH) of control and *NudC* RNAi groups, stained with DAPI to visualize DNA. Scale bar: 50 µm. (D) Quantification of PG size based on the area of a randomly selected lobe from control and NudC RNAi larvae. Each point represents one lobe; bars indicate mean ± SD (n=6). \*\**p* < 0.01, Mann–Whitney test. (E) Chromosomal morphology in single PG cells from control and *NudC* RNAi larvae. Scale bar: 2 µm. (F) DNA content (C-value) in control and *NudC* RNAi PG cells from wandering L3 larvae (94 hAH). Each point represents an individual cell; bars show mean ± SD (n=20). \*\*\*\**p* < 0.0001, Mann–Whitney test.

Taken together, these results indicate that NudC is required for maintaining proper cellular and chromosomal architecture in the PG. However, because cell size, microtubule organization, and polytene chromosome structure could be regulated not only in the PG but also in other polyploid cells, these findings may reflect a secondary effect related to compromised cellular integrity rather than the direct role of NudC in ecdysteroid biosynthesis.

### NudC ensures cell growth and tissue function in the salivary gland

Building on our findings and hypothesis above, we investigated whether *NudC* knockdown in other polyploid cells produces similar phenotypes. We focused on the larval salivary gland (SG), where cells grow large by undergoing endoreplication, achieving the C-value up to 1024 by the ninth replication cycle (Rodman, 1967). This makes the SG ideal for high-resolution imaging and detailed cellular analysis. Importantly, the SG is not essential for larval survival and growth, allowing gene manipulations without causing systemic effects.

We confirmed *NudC* expression in SG cells using NudC::GFP, a GFP-tagged endogenous NudC knock-in strain (Figure 2A). SG-specific knockdown of *NudC* via the *fkh-GAL4* driver efficiently reduced NudC::GFP levels (Figures 2A and 2B), and the RNAi larvae developed normally overall (Figure 2D). However, *NudC* RNAi caused a significant 70% reduction in SG tissue size (Figures 2E and 2F) and a 68% decrease in nuclear size compared to controls (Figures 2G and 2H). These phenotypes were consistently observed using a second independent RNAi line (Figures S2A-2D), where *NudC* expression was similarly reduced (Figure S2E). *NudC*-deficient SG cells also exhibited abnormal microtubule organization, showing relatively parallel and uneven microtubule distribution compared to controls (Figure S1C). Conversely, overexpression of *NudC* did not affect cell or nuclear size despite western blotting analysis confirming protein overload (Figures S3), suggesting that NudC is necessary but not sufficient for SG cell growth.

**Figure 2.**
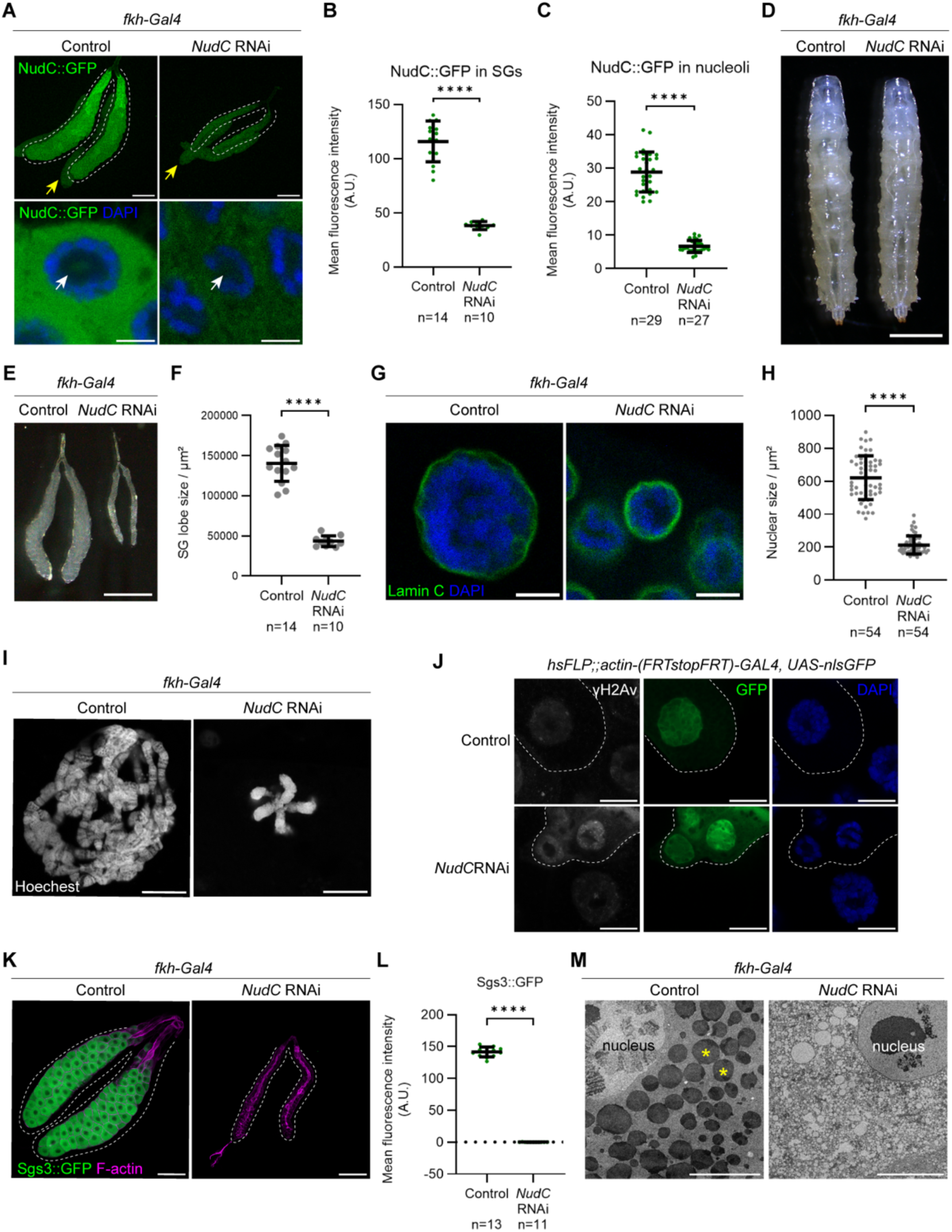
NudC is required for cellular growth, chromosomal integrity, and glue protein production in salivary gland (SG) cells. (A-C) NudC::GFP levels in control and *NudC* RNAi SGs from the L3 larvae. (A) NudC::GFP in control (*fkh>+, NudC::GFP*) and *NudC* RNAi (*fkh>NudC RNAi^THU^, NudC::GFP*) SGs. Associated fat bodies (indicated by yellow arrowheads) remained during dissection. Magnified views show SGs at 72 hAH, prior to the formation of large glue granules, allowing clear observation of NudC::GFP distribution. NudC::GFP was primarily cytoplasmic but also enriched in the nucleolus (white arrowheads). In *NudC* RNAi SGs, nucleolar NudC::GFP signals were markedly reduced. DNA was stained with DAPI (blue). Scale bars: 200 µm (upper panels); 10 µm (lower panels). (B) Quantification of mean fluorescence intensity (arbitrary unit, A.U.) of NudC::GFP in control (n=14) and *NudC* RNAi (n=10) SGs. (C) Quantification of nucleolar NudC::GFP fluorescence intensity in control (n=29) and *NudC* RNAi (n=27) SGs. Bars indicate mean ± SD. \*\*\*\**p* < 0.0001, Mann–Whitney test. (D) Representative wandering L3 larvae of control and *NudC* RNAi. Scale bar: 1 mm. (E and F) SG size in control and *NudC* RNAi larvae at the wandering L3 stage. (E) Images of control and *NudC* RNAi SGs. Scale bar: 0.5 mm. (F) Quantification of SG lobe area from control (n=14) and *NudC* RNAi (n=10) larvae. Bars show mean ± SD. \*\*\*\**p* < 0.0001, Mann–Whitney test. (G and H) Nuclear size in SG cells. (G) Representative nuclei labeled with Lamin C (green) and DAPI (blue). Scale bar: 10 µm. (H) Quantification of nuclear area from control (n=54) and *NudC* RNAi (n=54) SG cells. Nuclei were randomly selected from six cells in each of nine SGs per genotype. Bars show mean ± SD. \*\*\*\**p* < 0.0001, Mann–Whitney test. (I) Polytene chromosomes in control and *NudC* RNAi SG cells from the wandering L3 larvae, stained with Hoechst (white) and gently pressed to spread the chromosome arms. Scale bar: 10 µm. (J) Distribution of γH2Av (white) in SG mosaic clones from control and *NudC* RNAi larvae. Clones are marked by GFP and outlined with dashed lines; DNA is counterstained with DAPI (blue). Scale bar: 20 µm. (K and L) Glue protein production analyzed by Sgs3::GFP. (K) Representative SGs expressing Sgs3::GFP (green) with F-actin (magenta) under control (*fkh>+, Sgs3::GFP*) and *NudC* RNAi (*fkh>NudC RNAi^THU^, Sgs3::GFP*) conditions. Scale bar: 200 µm. (L) Quantification of Sgs3::GFP mean fluorescence intensity in control (n=13) and *NudC* RNAi (n=11) SGs. Bars represent mean ± SD. \*\*\*\**p* < 0.0001, Mann–Whitney test. (M) Transmission electron microscopy (TEM) images of control and *NudC* RNAi SG cells. Cytoplasmic glue granules (asterisks) abundant in control SG were scarcely observed in *NudC* RNAi SGs. Scale bar: 10 µm.

We further examined the impact of *NudC* knockdown on SG polytene chromosomes. Similar to the PG phenotype, *NudC* RNAi SG cells exhibited smaller polytene chromosomes characterized by the typical five-chromosome arm pattern (Figure 2I), implicating *NudC* in endoreplication control within SG cells. *NudC* knockdown also led to chromosome destabilization and increased DNA damage, evidenced by elevated levels of γH2Av, a marker of DNA double-strand breaks (Figure 2J).

Finally, we assessed SG functionality by analyzing glue protein secretion, which is essential for attaching the pupal case to a solid substrate during metamorphosis (Csizmadia et al., 2022). *NudC* RNAi SGs showed pronounced reduction of Sgs3-GFP fluorescence, a reporter for glue protein expression, (Figures 2K and 2L), and transmission electron microscopy (TEM) revealed a complete absence of secretory glue granules in *NudC* RNAi SG cells (Figure 2M). These findings demonstrate that NudC is critical for the major function of SGs.

### NudC regulates the nuclear and microtubule structure of larval polyploid cells

Analysis of the FLYATLAS 2 database (Krause et al., 2022) confirms that *NudC* is expressed in nearly all tissues, including polyploid SGs and fat bodies (see FlyBase FBgn0021768). Consistent with this, NudC::GFP strains showed NudC localization in multiple polyploid cells, such as the PG, epidermis, fat body, and also in diploid wing disc cells (Figure S4). Further supporting these expression patterns, *NudC*-deficient epidermal cells displayed disrupted microtubule organization (Figure S1E). These cells also showed altered polytene chromosome structure, accompanied by reduced nuclear size (Figure S1F). A comparable chromosomal phenotype was observed in fat body cells, although the high lipid content of this tissue prevented a clear assessment of microtubule organization (Figures S1G and S1H). These findings indicate that NudC is generally essential for maintaining cellular and chromosomal architecture in larval polyploid cells.

To further assess NudC function, we analyzed two *NudC* transheterozygous mutants (*NudC^GS15156^* / *NudC^Df(3L)ED223^* and *NudC^GS15156^* / *NudC^Df(3L)ED4674^*). These mutants exhibited significantly smaller larval body size, developmental delays, and lethality compared to the control *w^1118^* (Figure S5). In contrast, *fkh-GAL4*-driven *NudC* RNAi did not alter overall larval body size but strongly affected both SG size and morphology (Figure 2). As SG cells are substantially larger than those of the PG, they provide a robust model for studying NudC roles in polyploid cells. Based on these results, subsequent analyses focused on *fkh-GAL4*-driven *NudC* RNAi larvae to define the gene’s function in polyploid cell contexts.

### NudC localizes to both the cytoplasm and nucleolus in SGs

To clarify NudC function in SG cells, we examined the subcellular localization of NudC::GFP signal with greater detail. NudC was predominantly localized in the cytoplasm, but was also faintly, yet significantly, present in the nucleolus of SG cells (Figure 2A). This nucleolar signal is unlikely to be background noise, as it was markedly reduced upon *NudC* RNAi (Figures 2A and 2C). These observations suggest that NudC may participate in nucleolus-associated processes, such as ribosome biogenesis (RiBi), a process essential for protein synthesis.

### SG-specific *NudC* knockdown disrupts ribosome distribution and function, as well as associated cellular processes

To investigate the role of NudC in RiBi within SG cells, we performed ultrastructural analysis focusing on cytoplasmic ribosomes and rough endoplasmic reticulum (rER), the latter being a key site of protein synthesis marked by ribosomes attached to its cytosolic surface. In *NudC* RNAi SG cells, typical ribosome particles were rarely observed, and rER structures were largely absent (Figure 3A). This profound loss of ribosomes and rER prompted us to assess protein translation by labeling nascent proteins with O-propargyl puromycin (OPP), a direct indicator of ribosomal function. Compared to robust protein synthesis in control SGs, *NudC*-deficient cells exhibited a significant reduction in translational activity (Figures 3B and 3C).

**Figure 3.**
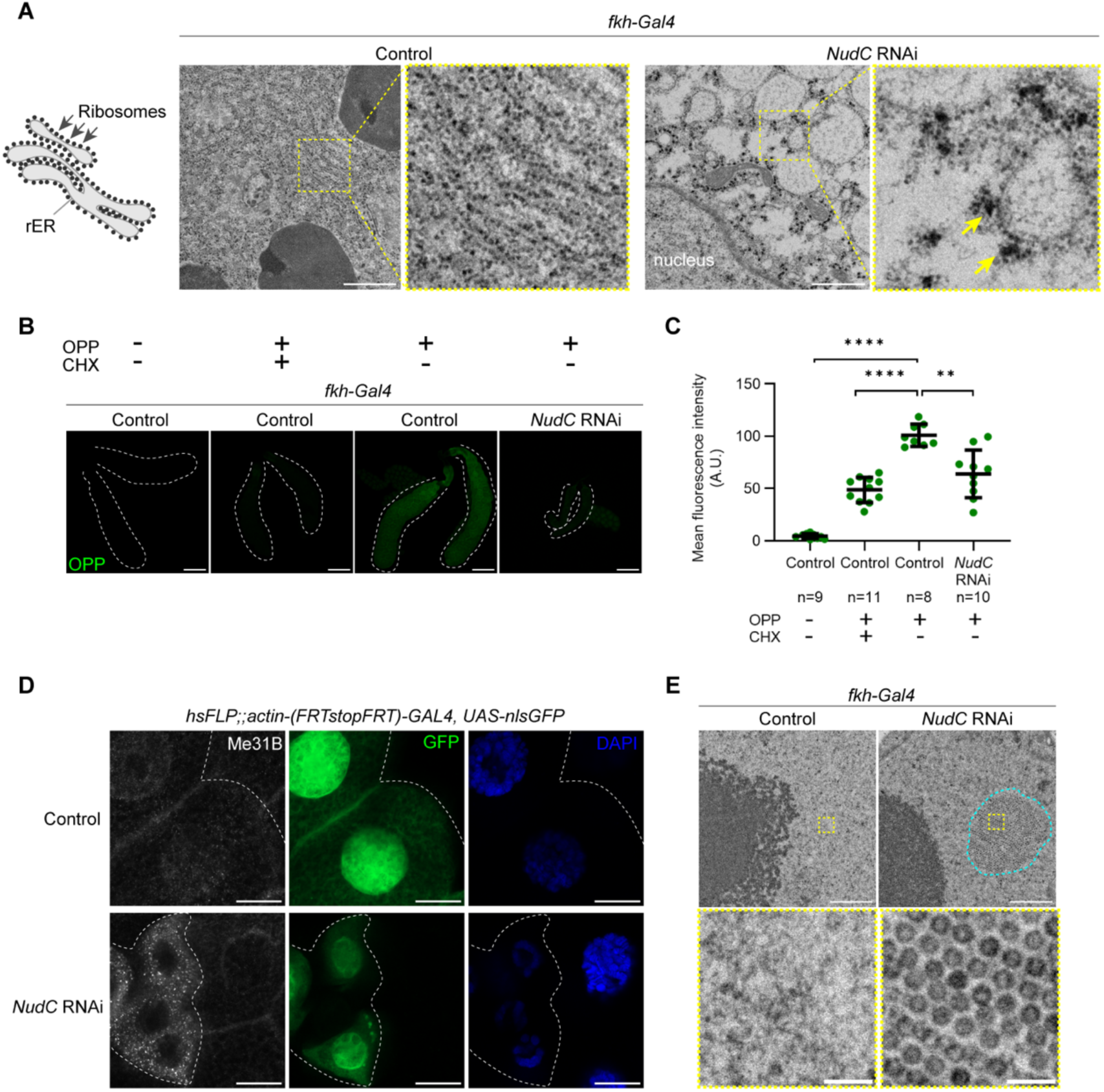
SG-specific *NudC* knockdown disrupts rER organization, ribosomal function, and associated cellular processes. (A) TEM images of cytoplasm from control and *NudC* RNAi SG cells at the wandering L3 stage. Enlarged views of boxed regions are shown to the right. In control cells, rough endoplasmic reticulum (rER) was readily detected, whereas it was largely absent in *NudC* RNAi cells. Arrowheads indicate electron-dense granules resembling processing (P)-bodies, which appeared larger than cytoplasmic ribosomes. Scale bars: 1 µm, for the panels on the left side of each experimental group. (B and C) Protein synthesis activity measured by OPP signals in SGs from the wandering L3 larvae. (B) Representative images of OPP signals (green) in SGs (outlined with dashed lines). Scale bar: 200 µm. (C) Quantification of OPP fluorescence intensity under control and *NudC* RNAi conditions, with or without cycloheximide (CHX) pretreatment (n = 8–11 SGs/group). CHX significantly reduced OPP signals, confirming specificity. Samples incubated without OPP serve as an additional negative control. Bars represent mean ± SD. \*\**p* < 0.01, \*\*\*\**p* < 0.0001; Brown–Forsythe and Welch ANOVA with Dunnett’s T3 multiple comparisons. (D) Localization of the P-body marker Me31B (white) in SG mosaic clones. Control and *NudC* RNAi cells were identified by GFP (outlined with white dashed lines). DNA is labeled with DAPI (blue). Scale bar: 20 µm. (E) Clusters of virus-like particles (VLPs) were observed in the nucleoplasm of *NudC* RNAi SG cells but not in controls. Representative TEM images show nucleoplasm of control (left) and *NudC* RNAi cells (right). VLP cluster is indicated by blue dashed circle; boxed regions are magnified below. Scale bars: 1 µm (upper panels) and 0.1 µm (lower panels).

TEM analysis also revealed numerous cytoplasmic electron-dense granules in *NudC* RNAi SG cells (Figure 3A), which we hypothesized to be processing (P)-bodies—stress-induced aggregates of untranslated mRNAs (Luo et al., 2018; Teixeira et al., 2005). This was confirmed by immunostaining for the P-body marker Me31B, showing abundant P-bodies within the cytoplasm of *NudC*-deficient SG cells (Figure 3D). These findings are consistent with the known role of P-bodies in mRNA storage and decay during translational arrest, suggesting that ribosomal dysfunction triggers mRNA decay pathways in *NudC*-deficient cells.

Additionally, many arrays of electron-dense particles were observed in the nucleoplasm of *NudC* RNAi SG cells, contrasting with their sparse presence typically seen in wild-type nuclei (Figure 3E). These particles resemble virus-like particles (VLPs), and their nuclear accumulation is associated with cellular stress (Strand and McDonald, 1985; Yoshioka et al., 1990), including ribosomal stress conditions (He et al., 2015; Wang and DiMario, 2017).

Together, these results demonstrate that loss of NudC function disrupts ribosome distribution and function, impairs rER integrity, and induces ribosomal stress responses, suggesting NudC’s critical role in maintaining ribosome abundance and associated cellular homeostasis.

### NudC is critical for maintaining rRNA levels in polyploid cells

To ascertain whether *NudC* maintains ribosome abundance by mediating RiBi in SG cells, we quantified rRNA levels using fluorescent *in situ* hybridization (FISH) with antisense probes targeting ITS1, ITS2, 18S, and 28S sequences (Figure 4A). The 18S and 28S rRNAs are core components of the small and large ribosomal subunits, respectively, while ITS1 and ITS2 are internal transcribed spacers located between them (Gerstberger et al., 2017). In SG cells, ITS1- and ITS2-containing pre-rRNAs are synthesized in the nucleolus (Figures 4B and 4C). Both mean fluorescence intensity and signal area of these pre-rRNAs were significantly reduced in *NudC* RNAi SG cells, resulting in a marked decrease in integrated intensity (Figures 4D-4G). Furthermore, 18S and 28S rRNA probes exhibited strong cytoplasmic signals and weaker nuclear fluorescence (Figures 4H and 4I), consistent with mature rRNA distribution. Both 18S and 28S rRNA signals were significantly diminished in *NudC*-deficient SG cells (Figures 4J and 4K). This reduction was specific to SGs, as rRNA levels in adjacent fat body cells remained unchanged (Figure S6). Northern blotting analysis validated these findings, showing decreased pre-rRNA and 28S rRNA bands in *NudC* RNAi SGs compared to controls (Figure 4L). In addition, *NudC* knockdown in fat body cells also led to decreased 18S and 28S rRNA levels, while ITS1 and ITS2 levels increased, indicating dysregulation of rRNA processing (Figure S7). These results collectively suggest that NudC is critical for maintaining rRNA levels and proper RiBi in polyploid cells.

**Figure 4.**
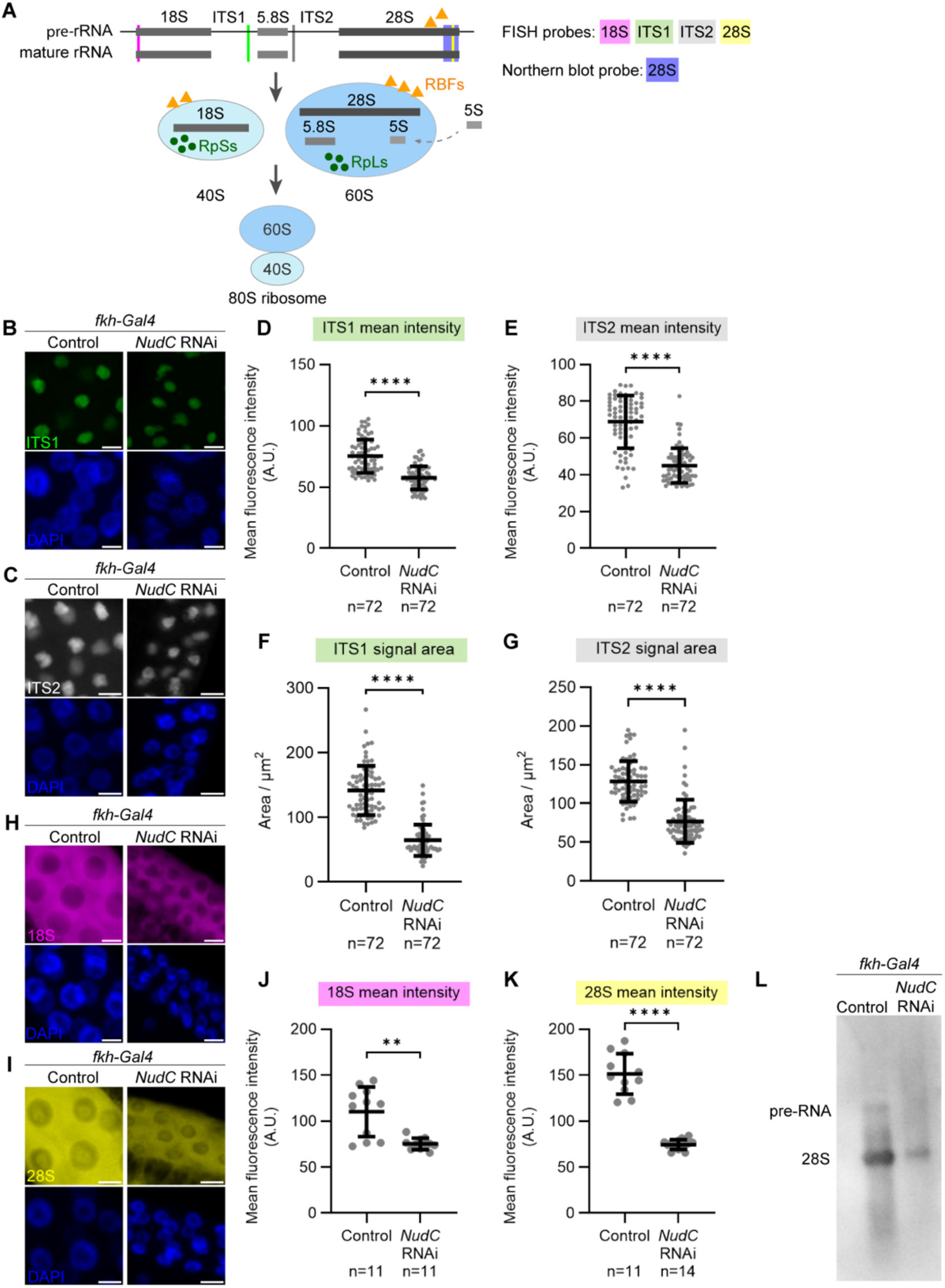
*NudC* is required to maintain rRNA levels in SGs. (A) Schematic representation of ribosome biogenesis (RiBi). Precursor rRNAs are synthesized in the nucleolus and cleaved at ITS1 and ITS2 sites to generate mature 18S, 5.8S, and 28S rRNAs. These assemble with ribosomal proteins (RpSs and RpLs) and 5S rRNA to form the 40S and 60S subunits, which combine to produce the functional 80S ribosome. This multi-step process is assisted by ribosome biogenesis factors (RBFs). (B-K) RNA fluorescence *in situ* hybridization (RNA FISH) for ITS1, ITS2, 18S, and 28S rRNAs in SGs from L3 larvae at 72 hAH. Probe regions are indicated in (A). Representative images show ITS1 (green), ITS2 (white), 18S (magenta), and 28S (yellow), with DNA stained by DAPI (blue). Scale bar: 20μm. (D–G) Quantification of ITS1 and ITS2 signals. Both mean fluorescence intensity and signal area were measured per cell. For each genotype, 72 cells were analyzed (six randomly selected from each of 12 SGs). (D and F) ITS1; (E and G) ITS2. (J and K) Quantification of mature rRNA levels. Mean fluorescence intensity of 18S (J) and 28S (K) rRNAs was measured in one randomly selected SG lobe per animal (n = 11–14). Bars represent mean ± SD. Mann–Whitney test: \*\**p* < 0.01, \*\*\*\**p* < 0.0001. (L) Northern blotting of control and *NudC* RNAi SG RNA from wandering L3 larvae using a probe against 28S rRNA (probe region indicated in A). Pre-rRNA and mature 28S bands were distinguished with an RNA size marker.

### Upregulation of genes encoding ribosomal proteins and ribosomal biogenesis factors in *NudC* knockdown SGs

RiBi is a highly regulated process that coordinates the assembly of rRNA and ribosomal proteins (RPs), guided by numerous RBFs (Figure 4A). Given the observed reductions in rRNA levels in *NudC* knockdown SG cells, we investigated whether RP and RBF gene expression was coordinately regulated in response to defective rRNA processing.

RNA sequencing (RNA-seq) analysis revealed widespread gene expression changes, with 2,089 genes down-regulated and 2,200 up-regulated in *NudC* RNAi compared to controls (Figure 5A). Gene Ontology (GO) analysis of the up-regulated genes showed enrichment of terms directly related to RiBi and rRNA processing within the top ten biological process categories (Figure 5B). Notably, cellular component categories such as nucleolus, ribosome, and preribosome featured prominently (Figure 5B). In contrast, the top down-regulated gene categories did not involve RiBi (Figures S8A and S8B). Consistent with this, most RP genes were up-regulated in *NudC*-deficient cells (Figure 5A). One up-regulated RBF was fibrillarin (Fib), a key player in pre-rRNA processing (Ojha et al., 2020). Fib mRNA upregulation was reflected by increased Fib protein levels in *NudC* knockdown SGs (Figures 5C and 5D), with accumulation observed both in the nucleolus and nucleoplasm. Similarly, the ribosomal protein RpS6 showed elevated protein levels in both the cytoplasm and nucleolus of *NudC*-deficient SG cells (Figures 5E and 5F).

**Figure 5.**
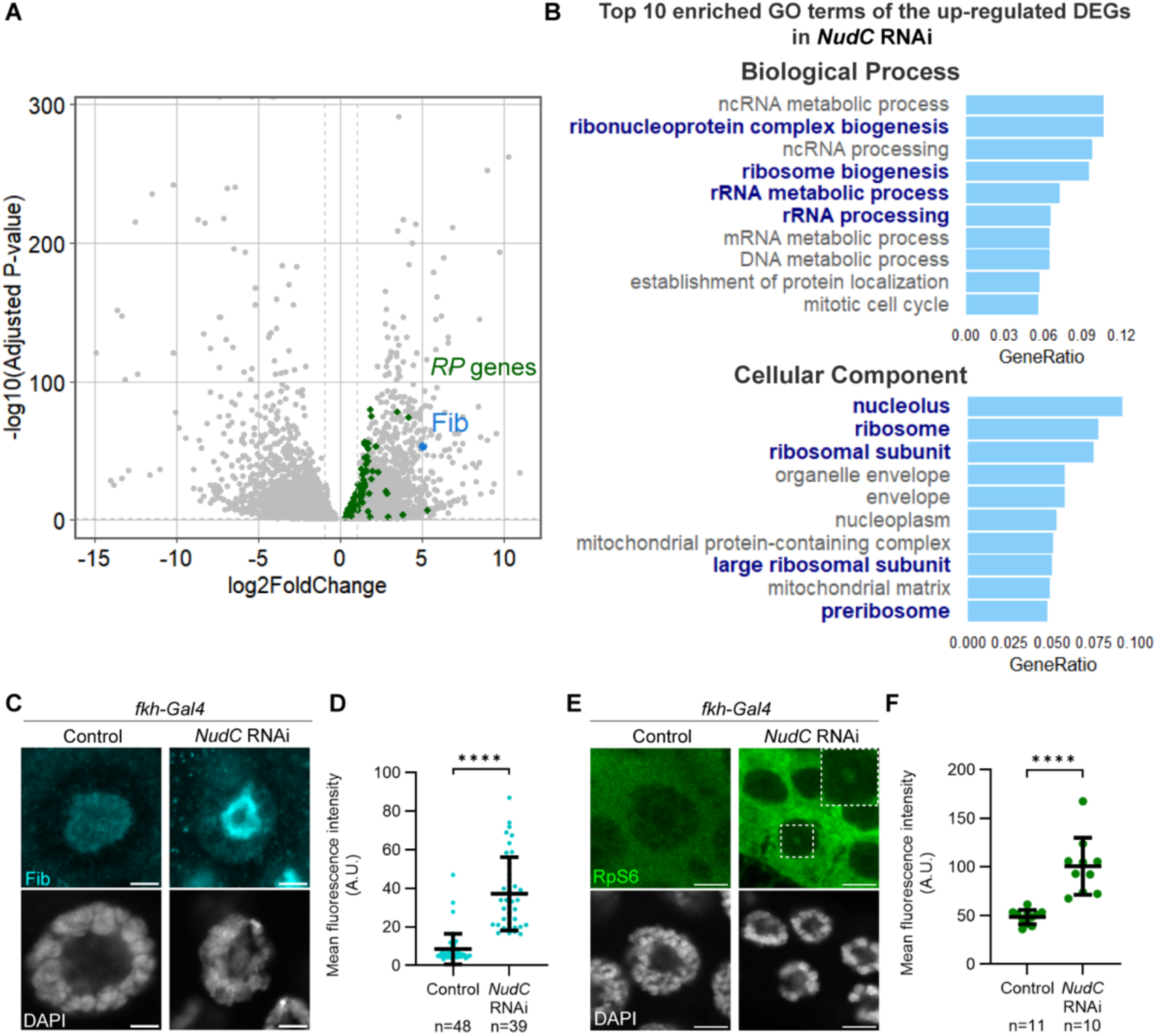
*NudC* knockdown in SGs induces an increase in the transcription and translation of RP and RBF genes. (A) Volcano plot showing differentially expressed genes (DEGs) between control and *NudC* RNAi SGs from wandering L3 larvae. Genes encoding RPs are highlighted in green; *fibrillarin* (*Fib*) is shown in blue. (B) Top 10 Gene Ontology (GO) enrichment terms (biological process and cellular component) among up-regulated DEGs (*p* < 0.05) in *NudC* RNAi SGs. Bars represent the proportion of up-regulated DEGs (“GeneRatio”). GO categories related to RiBi and nucleolus are emphasized in bold dark blue. (C and D) Fib localization and quantification. (C) Representative images of Fib (cyan) with DNA (DAPI, white) in control and *NudC* RNAi SG cells at the wandering L3 stage. Scale bar: 5 µm. (D) Quantification of Fib signal intensity (control: n = 48 cells; *NudC* RNAi: n = 39). Bars represent mean ± SD. \*\*\*\**p* < 0.0001, Mann–Whitney test. (E and F) RpS6 localization and quantification. (E) Representative images showing RpS6 (green) with DNA (DAPI, white) in control and *NudC* RNAi SG cells. Enlarged views of boxed regions (upper right) highlight increased RpS6 protein levels within the nucleolus of *NudC* RNAi cells. Scale bar: 10 µm. (F) Quantification of RpS6 signal intensity (control: n = 11 SGs; NudC RNAi: n = 10). Bars represent mean ± SD. \*\*\*\**p* < 0.0001, Mann–Whitney test.

Together, these data demonstrate that loss of *NudC* leads to decreased mature rRNAs and fewer ribosomes but an opposing upregulation of RPs and RBFs. This underscores the critical role of *NudC* in coordinating RiBi balance in SG cells.

### Functional overlap of NudC and RBFs in chromosome organization in SGs

Our previous findings led us to investigate whether knockdown of any RP or RBF gene would reproduce the distinct chromosomal abnormalities, such as the “five-blob” structure, observed in *NudC* knockdown polyploid cells. To address this, we re-evaluated data from the previous RNAi screen targeting PG cells (Ohhara et al., 2019). This re-evaluation revealed that DNA-enriched “blob” chromosomal structure was observed not only in *NudC* knockdown (Figure 1E), but also in 67 additional RNAi lines (Figure S9 and Table S1). To understand the biological functions of these genes, we conducted GO enrichment analysis, which showed that 40 of these genes encode RBFs (Table S2).

Building on this, we examined whether RNAi of RBF genes identified from the PG screen phenocopied *NudC* RNAi effects in SG cells. We focused on *eIF5* and *RpLP0-like*, whose roles of RiBi in *D. melanogaster* remain poorly defined, though their human orthologues, EIF5A and MRTO4, participate in pre-rRNA processing—a pathway regulated by NudC here (Tafforeau et al., 2013). We also included *Nopp140*, a well-characterized RBF involved in rRNA methylation in *D. melanogaster* (He et al., 2015), known to mediate chromosome structure in polyploid cells, including SGs (Cui and DiMario, 2007; He et al., 2015). RNAi of *eIF5*, *RpLP0-like*, and *Nopp140* in SG cells resulted in severely shortened polytene chromosome arms with disrupted banding patterns, mirroring *NudC* RNAi phenotypes (Figure 6A). Additionally, depletion of any of these genes elevated γH2Av levels, indicative of DNA damage response, similar to *NudC* knockdown (Figures 6B and 6C).

**Figure 6.**
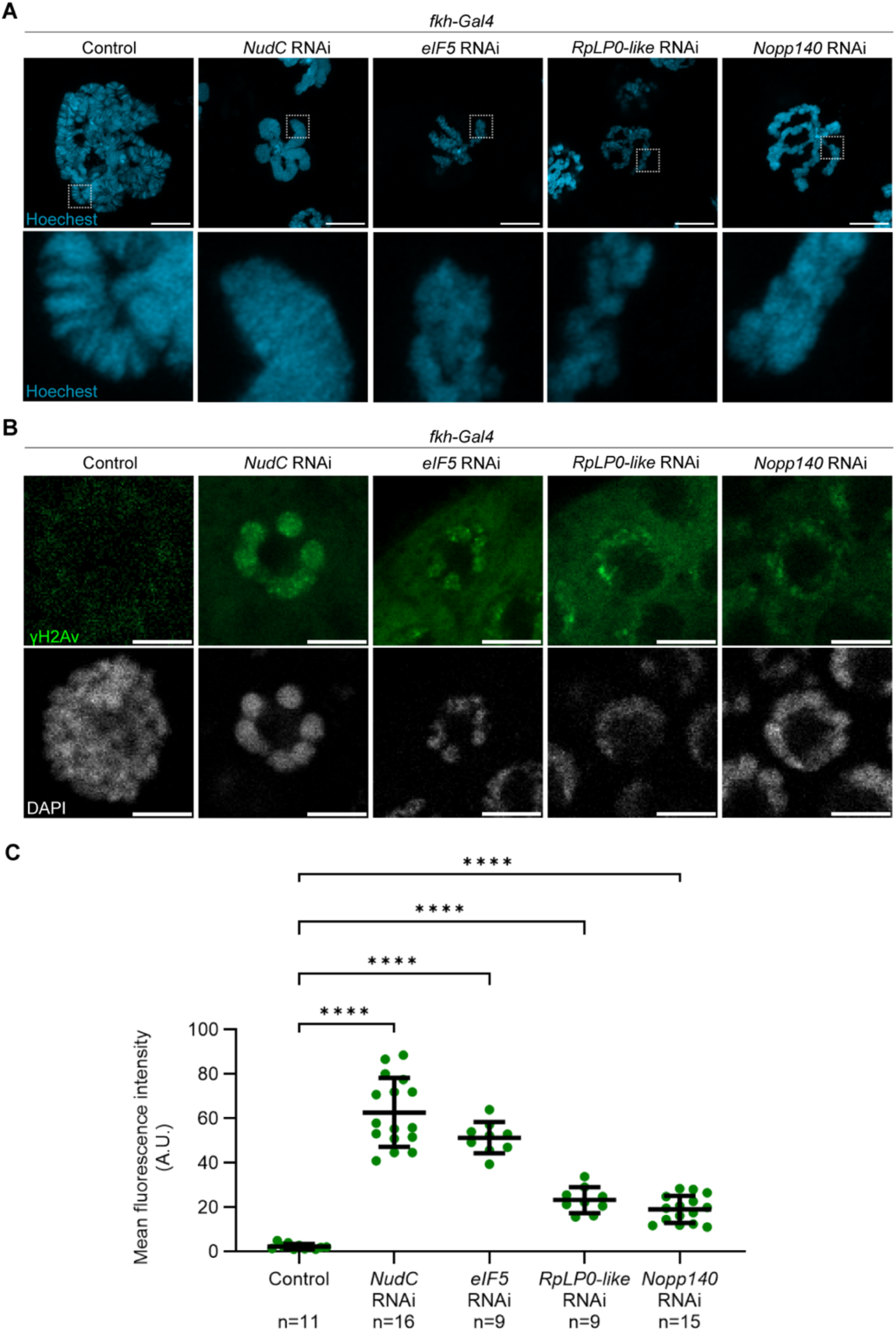
*NudC* knockdown phenocopies RBF loss by inducing chromosomal defects in SGs. (A) Polytene chromosomes from SG cells of wandering L3 larvae. Genotypes include control, *NudC* RNAi*, eIF5* RNAi, *RpLP0-like* RNAi, and *Nopp140* RNAi. Chromosomes were stained with Hoechst (cyan) and gently pressed to spread the arms. Boxed regions are shown at higher magnification below. Scale bar: 10 µm. (B and C) DNA damage in SGs assessed by γH2Av staining. (B) Representative images of γH2Av (green) with DNA counterstained by DAPI (white). Scale bar: 10 µm. (C) Quantification of γH2Av fluorescence intensity in SGs from control, *NudC* RNAi*, eIF5* RNAi, *RpLP0-like* RNAi, and *Nopp140* RNAi larvae (n = 9–16 SGs/group). Bars show mean ± SD. \*\*\*\**p* < 0.0001 (Brown–Forsythe and Welch ANOVA, followed by Dunnett’s T3 multiple comparisons).

These results collectively underscore the role of NudC alongside RBFs in maintaining chromosomal integrity in *D. melanogaster* SG cells.

### Knockdown of one RBF induces the upregulation of RPs and other RBFs in SGs

We hypothesized that the increased expression of RP and RBF genes in *NudC* RNAi SG cells reflects a compensatory response aimed at restoring ribosomal function. If this mechanism holds, loss-of-function in any RP or RBF gene may similarly induce a compensatory response, by which most RP and RBF genes become transcriptionally up-regulated.

To test this, we performed RNA-seq on SGs with RNAi of *eIF5*, *RpLP0-like*, and *Nopp140*. Knockdown of each gene caused extensive transcriptional changes, with the majority of RP genes being up-regulated (Figures 7A-7C). GO analysis of up-regulated genes confirmed enrichment for ribosome or rRNA processing, paralleling the response observed in *NudC* RNAi cells (Figures 7D-7I). Down-regulated gene sets showed no enrichment for RiBi-related pathways (Figures S8C-8H), consistent with a specific compensatory activation.

**Figure 7.**
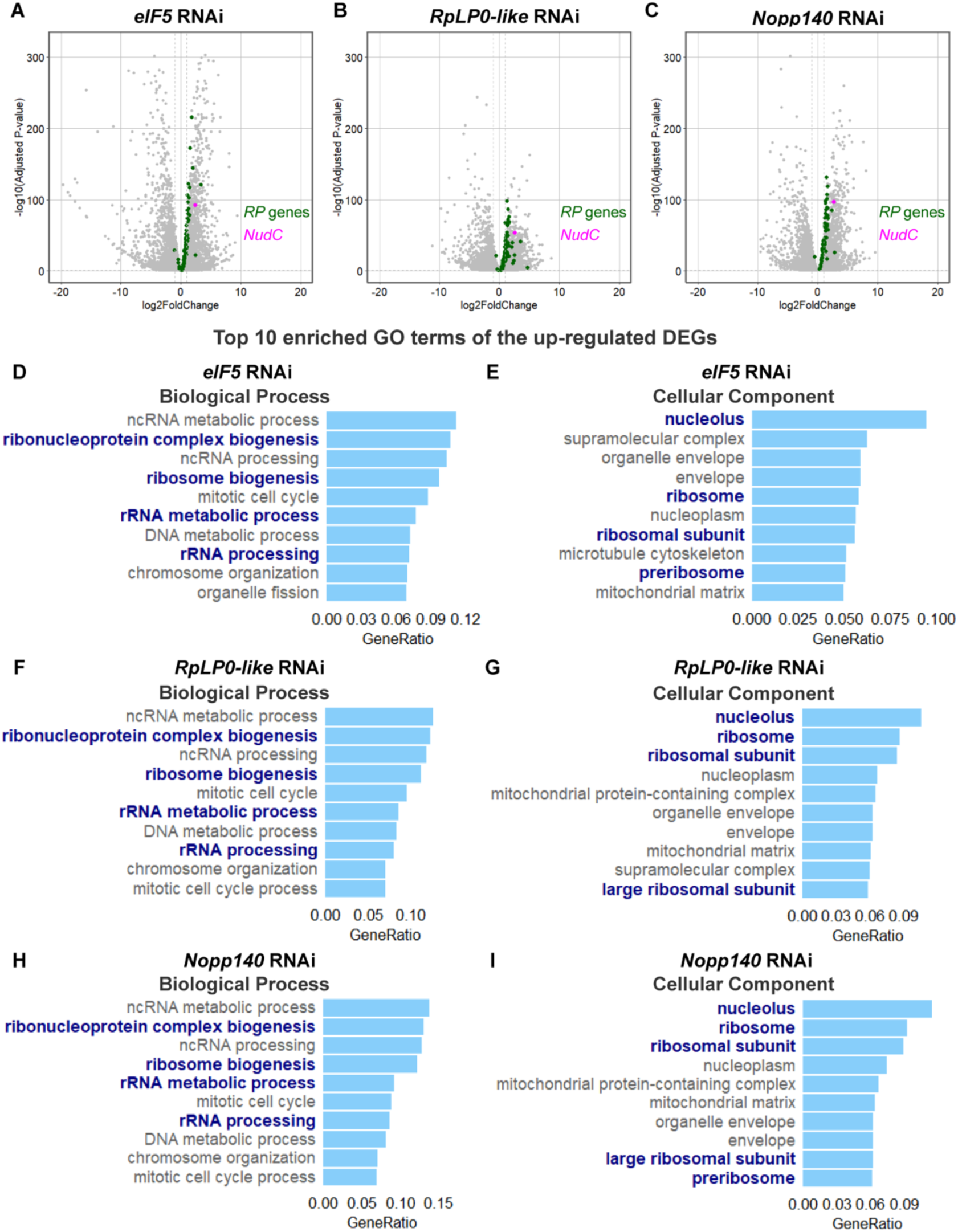
Knockdown of RBF genes upregulates *NudC* and RiBi-related genes in SGs. (A-C) Volcano plots showing DEGs in SGs from wandering L3 larvae upon *eIF5* RNAi, *RpLP0-like* RNAi, and *Nopp140* RNAi. RP genes are highlighted in green, and *NudC* is shown in magenta. (D-I) GO enrichment of up-regulated DEGs (*p*<0.05) in the same conditions. The top 10 enriched terms are shown for biological process and cellular component categories. Bars represent the proportion of up-regulated DEGs (“GeneRatio”). GO categories related to RiBi and nucleolus are emphasized in bold dark blue.

Intriguingly, *NudC* expression was strongly elevated following depletion of these RBFs (Figures 7A-7C), emphasizing its pivotal role in responding to impaired RiBi and coordinating compensation to maintain ribosomal homeostasis. These findings suggest a robust cellular mechanism where disruption of RiBi triggers widespread transcriptional feedback to preserve ribosomal function.

### *NudC* knockdown induces stress responses associated with impaired RiBi in SGs

Previous studies have shown that loss of RBFs activates the stress-responsive c-Jun N-terminal kinase (JNK) pathway and induces autophagy in polyploid cells (DeLeo et al., 2021; James et al., 2013; Rosby et al., 2009; Wang and DiMario, 2017). Consistent with these reports, *NudC* knockdown in SG cells led to a significant increase in phosphorylated JNK levels (Figure 8A). There was also a pronounced accumulation of Atg8a and acidic autolysosomes, indicating enhanced autophagy (Figures 8B and 8C). Notably, *NudC* knockdown did not result in apoptosis (Figure S10). Together, these results reinforce the role of NudC in regulating the stress response induced by ribosome disruption in polyploid cells.

**Figure 8.**
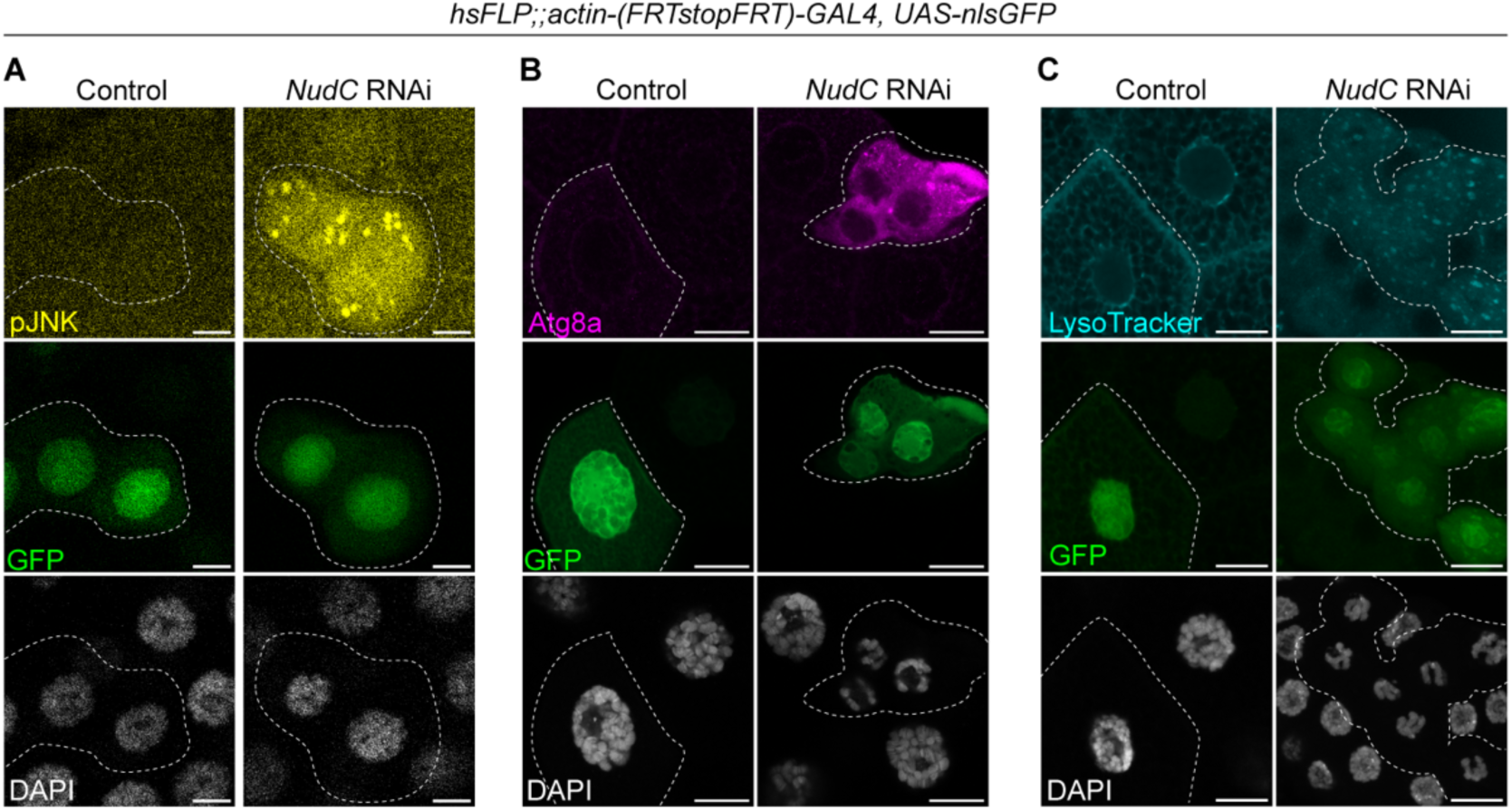
*NudC* knockdown activates JNK signaling and induces autophagy in SGs. Localization of phosphorylated-JNK (pJNK, A), Atg8a (B), and acidic lysosomes (LysoTracker, C) in control and *NudC* RNAi SG clones at the wandering L3 stage. Clones are marked by GFP (outlined with dashed lines). DNA is counterstained with DAPI (white). Scale bar: 20 µm.

### The role of NudC in RiBi might be dynein-independent

NudC is known to interact with dynein and its regulator Lis1, regulating microtubule-mediated cellular events (Kawano et al., 2022; Riera and Lazo, 2009). To assess whether the role of NudC in RiBi within polyploid cells depends on its established involvement with dynein, we performed SG-specific RNAi of *dynein intermediate chain* (*Dic*), *dynein heavy chain* (*Dhc*), and *Lis1*. Depletion of *Dic* or *Lis1* did not affect SG size or chromatin structure (Figures S11A and S11B). In contrast, *Dhc* knockdown caused a substantial reduction in tissue size, yet polytene chromosome morphology remained unchanged (Figures S11A and S11B). Additionally, protein translation was not inhibited in *Dhc* depletion (Figures S11C and S11D). These findings suggest that the role of NudC in maintaining RiBi operates independently of its role in dynein-mediated microtubule dynamics.

## Discussion

Our findings demonstrate that NudC is indispensable for RiBi in the polyploid cells of *D. melanogaster* larvae. *NudC* knockdown affects multiple key cellular processes, including cell and nuclear growth, polytene chromosome organization, and autophagy. These defects likely result from *NudC* depletion, altering ribosome distribution and causing a significant reduction in rRNA levels, highlighting the critical role played by NudC in maintaining ribosome abundance. Moreover, knockdown of *NudC* led to an increase in the expression of genes encoding RPs and RBFs, likely reflecting a compensatory cellular response to reduced rRNA synthesis. This compensatory upregulation, along with similar chromosomal defects and stress responses following RBF depletion, suggests that NudC and RBFs participate in overlapping regulatory pathways that preserve ribosomal function and cellular homeostasis. Notably, our results indicate that the role of NudC in RiBi is independent of its established function in dynein regulation, supporting the existence of a distinct mechanism governing its involvement in RiBi.

Our results demonstrate that NudC regulates rRNA levels in polyploid SG and fat body cells. NudC::GFP localizes at detectable levels within the nucleolus of wild-type cells from both tissues, and *NudC* depletion significantly reduces nucleolar signal intensity, supporting a direct role for NudC in rRNA processing at this primary site of RiBi. The present study showed a dramatic increase in the expression of RP and RBF genes in polyploid cells with reduced rRNA synthesis. This upregulation was observed not only with *NudC* knockdown but also following RNAi of RBF genes, indicating a compensatory homeostatic mechanism that maintains stable RiBi in polyploid cells. To date, the homeostatic regulation of RiBi in *D. melanogaster* remains poorly investigated. Previous studies reported reduced rRNA levels alongside accumulation of certain RPs within nucleoli of SGs (Martins et al., 2017).

Similarly, depletion of either Nucleostemin 1 or 2, which are pre-rRNA processing factors, leads to the accumulation of RPs in the nucleolus of midgut polyploid cells (Rosby et al., 2009; Wang and DiMario, 2017). Our study provides a more comprehensive view, showing that impaired pre-rRNA processing upregulates most RP genes and causes accumulation of RPs in both the nucleolus and cytoplasm of polyploid cells. It should be noted that although depletion of pre-rRNA processing factors Nopp140 and Nucleostemin 2 led to decreased levels of ribosomal proteins RpL23a and RpL34 in whole *D. melanogaster* larval lysates (James et al., 2013; Wang and DiMario, 2017), we observed an upregulation of these genes specifically in *Nopp140*-depleted SGs. This difference suggests that compensatory or homeostatic mechanisms may differentially regulate RP abundance depending on the cellular context. Notably, loss of RP genes disrupts rRNA synthesis in both *D. melanogaster* diploid wing disc cells and human cultured cells (Ajore et al., 2017; Juli et al., 2016; Kiparaki et al., 2022). Whether RP-depletion similarly affects rRNA levels in polyploid cells of *D. melanogaster* warrants further investigation. These highlight the complexity of homeostatic regulation in RiBi and underscore the importance of analyzing distinct tissues to fully understand the molecular consequences of perturbing RBFs.

*NudC* knockdown exhibited reduced nuclear size and disrupted endoreplication. Given a general rule of a consistent nuclear-cytoplasmic ratio (Balachandra et al., 2022; Neurohr et al., 2019), the disrupted endoreplication in *NudC* knockdown polyploid cells may account for smaller tissue size. We also observed “five-blob” polytene chromosomes in *NudC* RNAi polyploid cells. Considering that a similar chromosome structure transiently appears in developing ovarian nurse cells (Baxley et al., 2011; Dej and Spradling, 1999), the observed phenotype suggests a developmental arrest in DNA replication. Alternatively, NudC may play a more direct role in shaping the tertiary structure of polytene chromosomes. Recent evidence further suggests that RBFs can impact chromosome architecture. For example, mutations in *Non3*, an orthologue of *ribosome production factor 2*, have been shown to affect the formation and function of the nucleolus, and suppress correct chromatin packaging in *D. melanogaster* (Yushkova et al., 2025). Pre-rRNA can also promote chromosome dispersion during mitosis (Ma et al., 2022), indicating the broader involvement of RBFs in chromosome structure regulation. However, how NudC and RBFs such as Non3 contribute to chromosome organization in post-mitotic cells requires further investigation.

Disrupting RiBi by knockdown of *NudC* triggers nucleolar stress responses in polyploid cells, including the formation of cytoplasmic P-bodies, dense clusters of VLPs in the nucleoplasm, DNA damage, autophagy, and activation of the JNK signaling pathway. These stress responses closely mirror those observed in *D. melanogaster* following loss of RBFs, such as *Nopp140* depletion (DeLeo et al., 2021; He et al., 2015; James et al., 2013). Nucleoplasmic VLPs are believed to originate from retrotransposons and may contribute to cellular adaptation and survival under stress conditions, including those involving ribosomal stress (He et al., 2015; Nelson et al., 2023; Wang et al., 2022; Wang and DiMario, 2017). Their presence in *NudC*-deficient cells likely represents a nucleolar stress response, reflecting the cell’s efforts to cope with disrupted RiBi. The nucleolus itself acts as a central sensor and integrator of cellular stress signals. Mechanistically, p53 mediates nucleolar stress responses in mammalian cells (Donati et al., 2011), whereas in *D. melanogaster*, this regulatory role is primarily governed by the JNK signaling pathway, which is strongly activated following RiBi disruption (DeLeo et al., 2021; James et al., 2013). Activation of JNK can promote autophagy, as this pathway positively regulates autophagy-related genes, particularly in larval midgut and fat body cells (Wu et al., 2009). This mechanism illustrates a similar adaptive response observed in *NudC*-deficient cells, where JNK signaling manages nucleolar stress by promoting autophagy. We observed increased γH2Av signals in SG cells following *NudC* knockdown, though the biological meaning of this phenomenon remains unclear. Notably, γH2Av activates poly(ADP-ribose) polymerase 1, a key regulator of precursor rRNA processing, post-transcriptional modification, and pre-ribosome assembly (Boamah et al., 2012; Kotova et al., 2011). This suggests that elevated γH2Av levels in *NudC*-depleted SGs represent a cellular response linked to RiBi disruption.

We also found that NudC influences RiBi in polyploid cells in a tissue-dependent manner. Loss of *NudC* reduces 18S and 28S rRNA levels in both SG and fat body cells, but the effects on ITS1 and ITS2 vary between these tissues. Since ribosome abundance generally correlates with protein synthesis, cells with high protein production, such as those found in SGs, may be especially sensitive to disruptions in RiBi. Additionally, chromosomal architecture defects caused by *NudC* RNAi vary among polyploid cell types. The “five-blob” phenotype is commonly observed in PG and epidermal cells, but less apparent in SG and fat body cells, possibly due to differences in polytene chromosome size (Zielke et al., 2013). Overall, these findings suggest that NudC maintains rRNA levels within the nucleolus and preserves chromosome structure, with its effects varying according to cell type-specific mechanisms.

Though it has been well established as necessary for microtubule-associated dynein regulation, our results indicate that NudC also plays a role in a distinct, dynein-independent function in RiBi. Although we observed that microtubule disorganization following *NudC* knockdown, targeting dynein components or *Lis1* did not replicate the reductions in cell and nuclear size, the alterations in polytene chromosome structure, and the decline in translational activity caused by *NudC* depletion in polyploid cells. Of note, the regulatory relationship between NudC and Lis1 varies by species: in zebrafish, Lis1 functions downstream of NudC (Kawano et al., 2022), whereas in *NudC* knockout mice, Lis1 levels remain unchanged (Garner et al., 2024). This suggests that the molecular mechanisms linking NudC and dynein may differ across species.

In summary, this study uncovers new moonlighting functions of NudC, highlighting its critical role in RiBi within polyploid cells. Given the essential contribution of RiBi to the development of various ribosomal diseases (Jiao et al., 2023), our findings may inform future research on the molecular bases of these disorders and open avenues for therapeutic development targeting ribosome homeostasis.

## Materials and Methods

### Key resources table

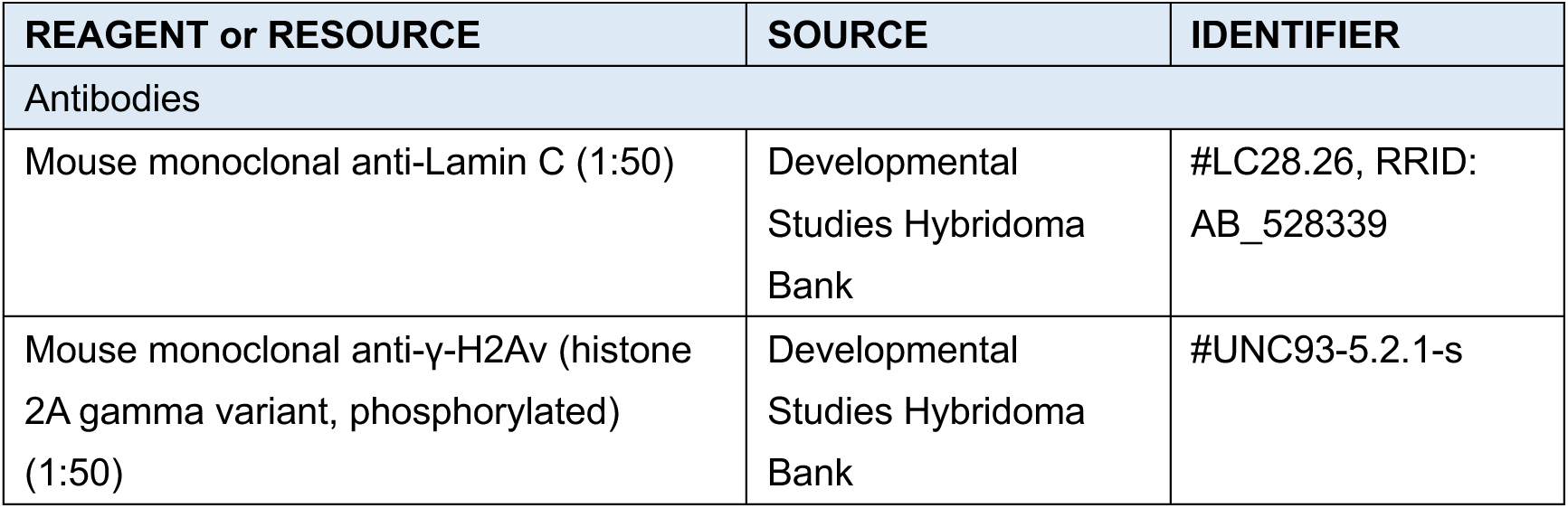
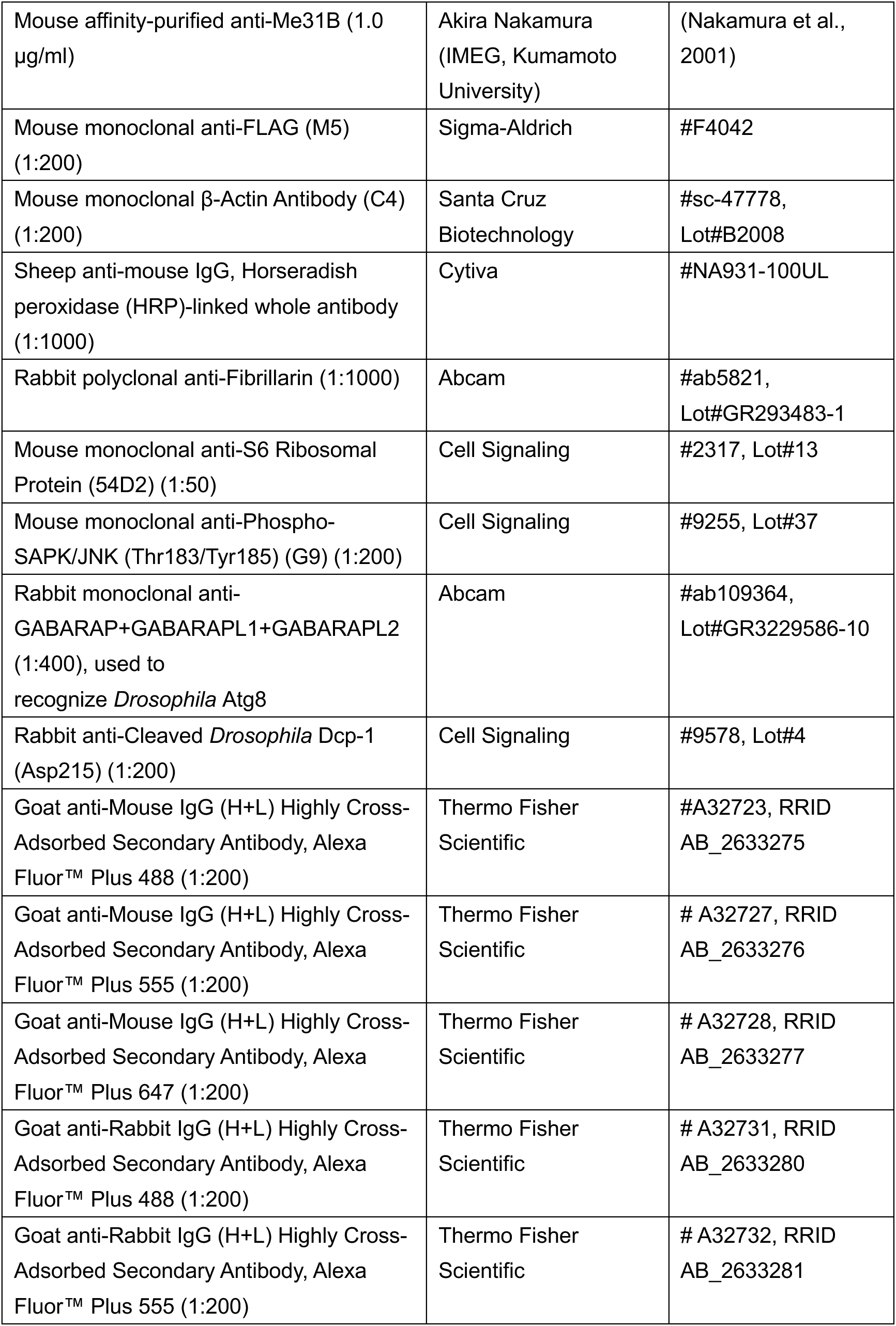
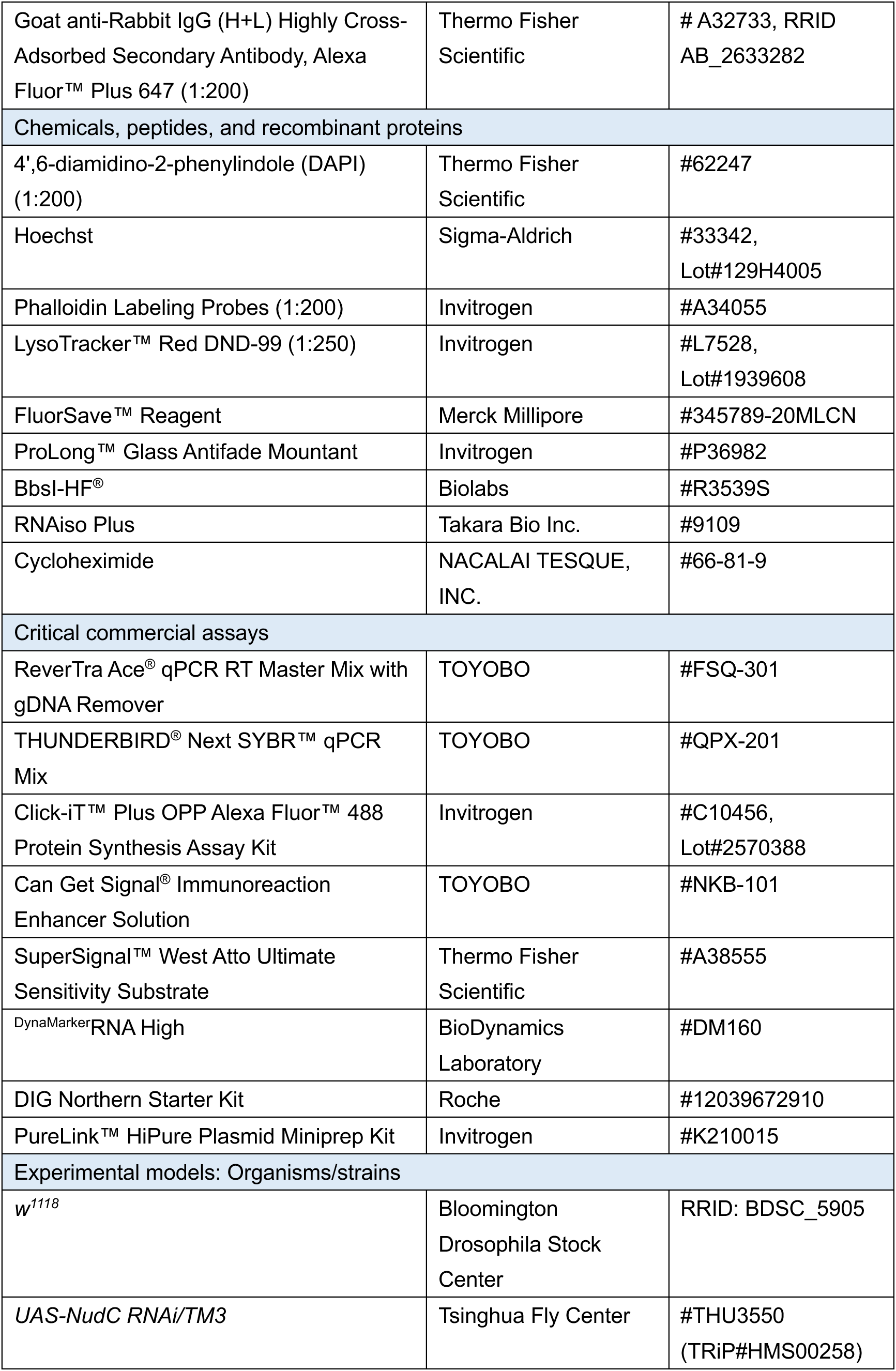
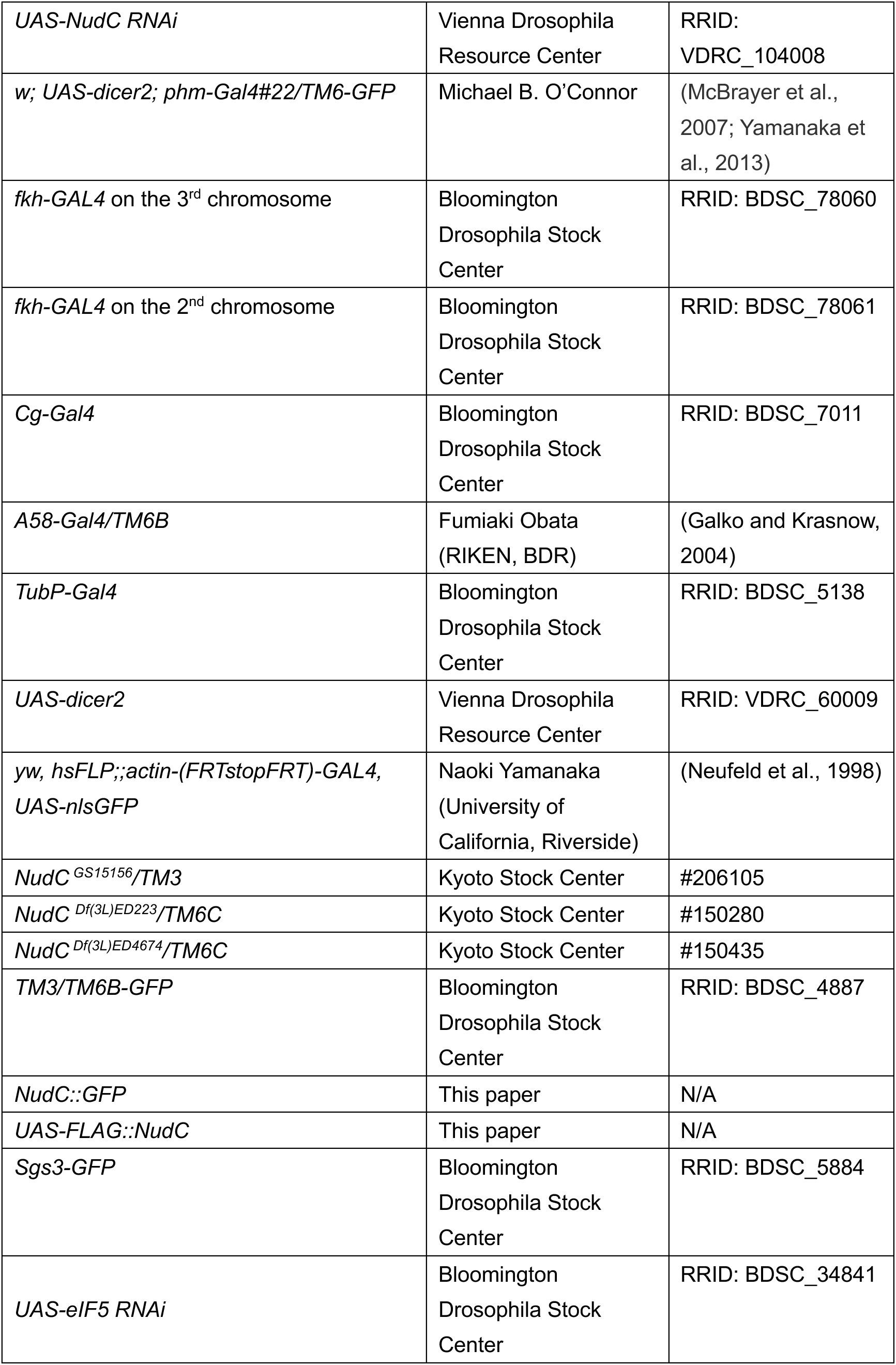
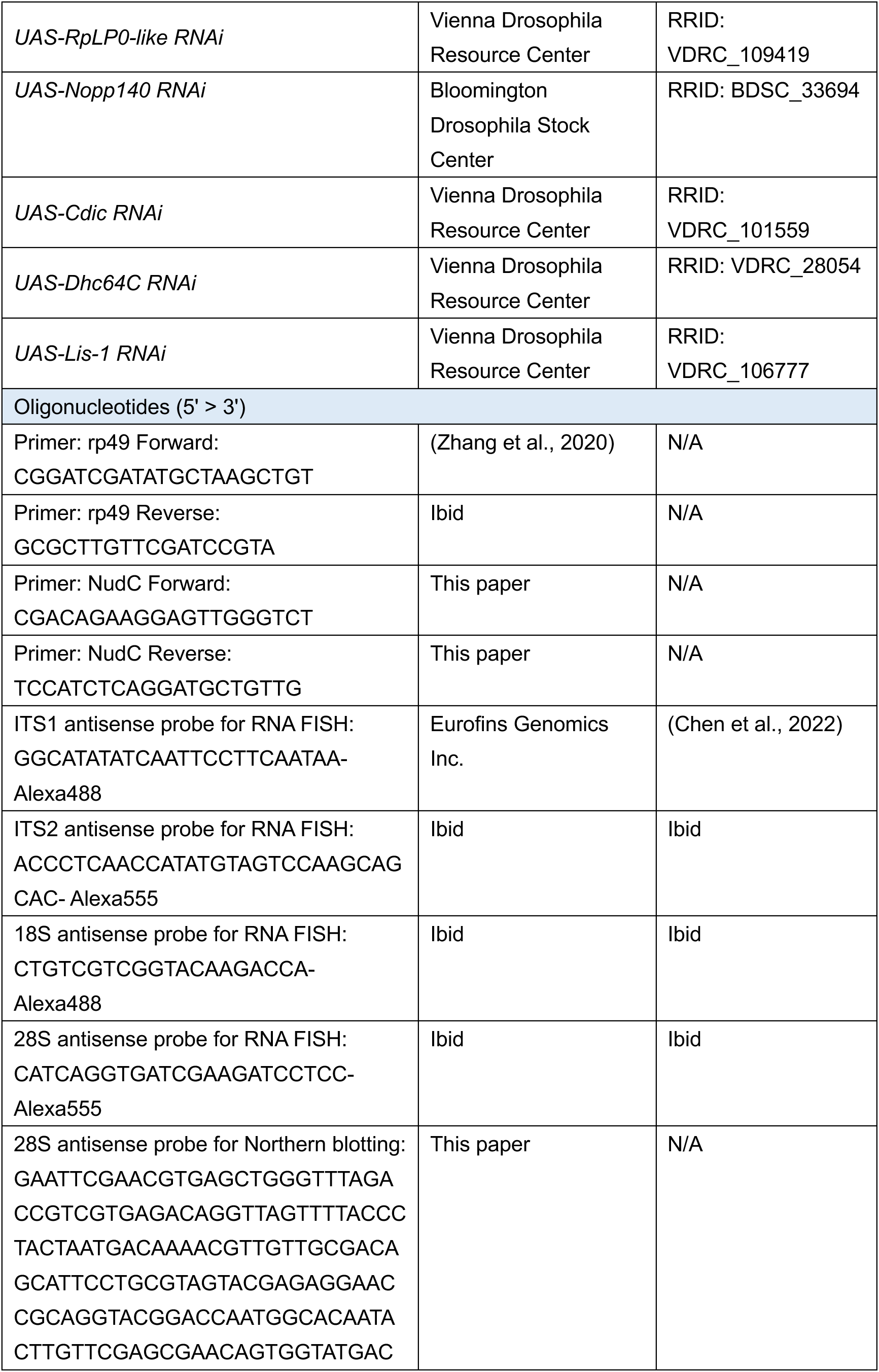
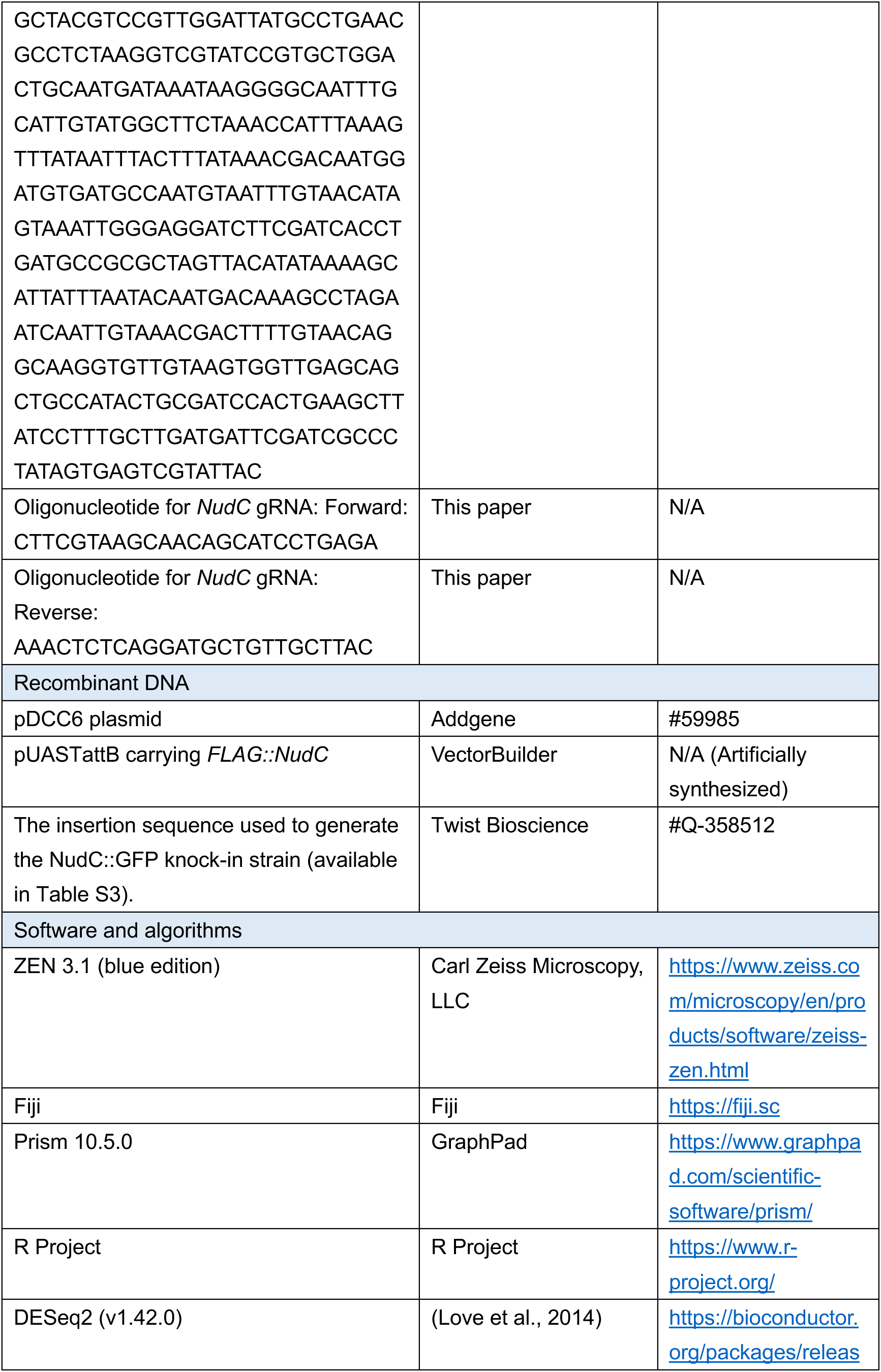
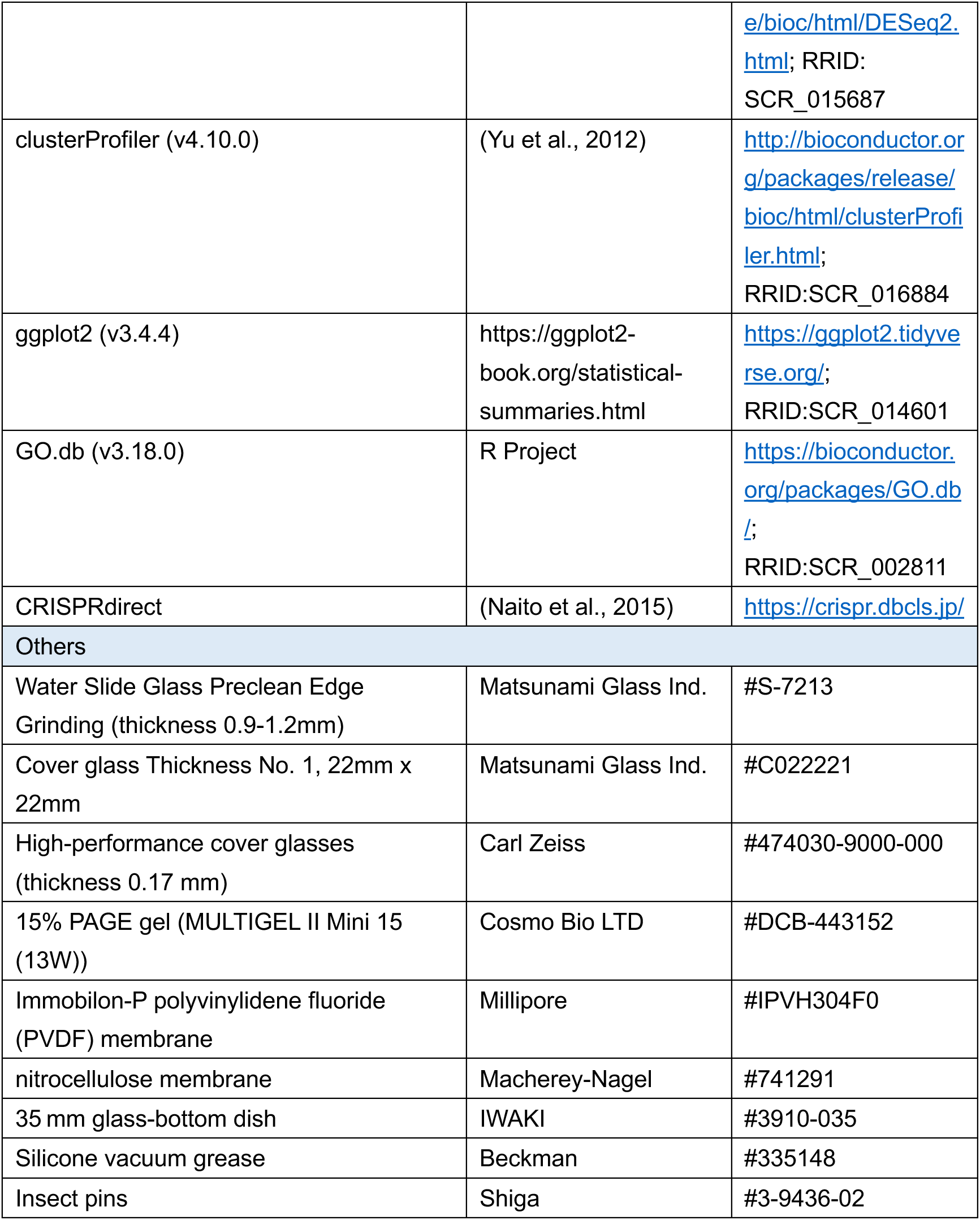

### Fly stocks and genetics crosses

*Drosophila melanogaster* flies were cultured on a standard diet (0.55 g agar, 10.0 g glucose, 9.0 g cornmeal, 4.0 g yeast extract, 300 μl propionic acid, 350 μl 10% butylparaben in 70% ethanol, and 100 ml water). Unless otherwise stated, experiments with flies were conducted at 25 °C under a 12:12-hour light/dark cycle. All fly strains used in the experiments are listed in the <u>Key Resources Table</u>. Larvae were not separated by sex except during fat body analysis, due to known sex-dependent differences in fat body cell size (Rideout et al., 2015). Fat bodies adjacent to the male larval gonad were specifically examined.

*UAS-NudC RNAi* (#THU3550), *NudC^GS15156^, NudC^Df(3L)ED223^,* and *NudC^Df(3L)ED4674^* strains were balanced with the *TM6-GFP* balancer. To generate the two *NudC* transheterozygous mutants, *NudC^GS15156^/TM6-GFP* flies were crossed with either *NudC^Df(3L)ED223^/TM6-GFP* or *NudC^Df(3L)ED4674^/TM6-GFP*.

### Salivary gland clone induction

Mosaic clones were generated by crossing *yw, hsFLP;;actin-(FRTstopFRT)-GAL4, UAS-nlsGFP* flies with either *w^1118^*or *UAS-NudC RNAi* (#THU3550). Embryos were incubated in small glass vials containing food for 1 hour after egg laying at 25 °C, then subjected to a 5-minute heat shock at 37 °C in a water bath. After the heat shock, embryos were returned to 25 °C. This method typically generated mosaic clusters within the SG, each consisting of several GFP-positive cells.

### Generation of transgenic flies

The NudC::GFP knock-in strain was generated using a CRISPR-Cas9-based approach, tagging the endogenous *NudC* at its C-terminus with the GFP coding sequence. Suitable guide RNA (gRNA) target sites within the *NudC* genomic region were identified using CRISPRdirect (Naito et al., 2015). The gRNA-expressing plasmid was constructed by annealing oligonucleotides (see <u>Key Resources Table</u>) and cloning them into a BbsI-digested pDCC6 vector. A pTwist-Amp-High-Copy-based plasmid containing two homology arms with the *GFP* coding sequence (Table S3) was obtained from Twist Bioscience (San Francisco, CA, USA). Plasmids were purified with PureLink™ HiPure Plasmid Miniprep Kit (Invitrogen, Waltham, MA, USA) and injected into *w^1118^* embryos for the knock-in integration. Successful integration of the NudC::GFP construct was confirmed by sequencing with the SeqStudio™ Genetic Analyzer System (Applied Biosystems, Waltham, MA, USA).

A pUASTattB-based plasmid carrying *NudC* with an N-terminal FLAG tag was synthesized by VectorBuilder, Inc. (Chicago, IL, USA), and the purified plasmid was injected into *w^1118^* embryos.

### Quantitative reverse transcription PCR (RT-qPCR)

RNA was extracted from control (*tubP>dicer2, +*) and *NudC* RNAi (*tubP>dicer2, NudC* RNAi) larvae at 40 hours after hatching (hAH) using RNAiso Plus reagent (Takara Bio Inc., Shiga, Japan). cDNA was synthesized with ReverTra Ace^®^ qPCR RT Master Mix with gDNA Remover (TOYOBO, Osaka, Japan). qPCR was performed on the Thermal Cycler Dice^®^ Real Time System TP800 (Takara Bio Inc.) using THUNDERBIRD^®^ Next SYBR™ qPCR Mix (TOYOBO). Target mRNA levels were normalized to *ribosomal protein 49* (*rp49*), and relative expression changes were calculated. Primer sequences used are listed in the <u>Key Resources Table</u>.

### Ploidy measurements

The C-value in the PG was measured following a previously described protocol (Ohhara et al., 2019) with modifications. Larvae at 94 hAH and adult male flies were dissected in 1× phosphate-buffered saline (1× PBS) (pH 7.4). The brain-ring gland complex from larvae and seminal vesicles from male flies fixed in 4% paraformaldehyde (PFA) in 1× PBS for 30 minutes. After fixation, tissues were washed thrice for 10 minutes in 0.3% PBT (0.3% Triton X-100 in 1× PBS) and once for 5 minutes in 1× PBS. Samples were then stained with DAPI (Thermo Fisher Scientific, Waltham, MA, USA) for 30 minutes, followed by three washes in 1× PBS for 5 minutes each. Tissues were mounted in FluorSave™ Reagent (Millipore, Burlington, MA, USA) and covered with a cover glass (Matsunami, Osaka, Japan). Seminal vesicles on each slide were imaged using consistent settings and gain to allow sperm cells to serve as internal controls. Images were acquired with a Zeiss LSM900 confocal microscope, and ZEN 3.1 was used for optimizing image acquisition, collecting z-stacks, and generating maximum intensity projections. DAPI intensity and area were quantified using Fiji/ImageJ (https://fiji.sc), with the integrated intensity (mean intensity × area) of sperm cells (1C) on the same slide set as 1. The integrated intensity of PG cells was then calculated relative to this standard to determine their C-value.

### Immunostaining

Larval tissues were dissected in 1× PBS and fixed with 4% PFA for 30 minutes at room temperature. Samples were washed thrice with 0.3% PBT and then blocked in 2% BSA in PBT for 1 hour at room temperature. Following blocking, tissues were incubated overnight at 4 °C with primary antibodies. After three washes with PBT, samples were incubated with secondary antibodies for 2 hours at room temperature. Details of all antibodies used are listed in the <u>Key Resources Table</u>. To visualize tissue or cell morphology, filamentous actin (F-actin) was stained using Phalloidin-Alexa-Fluor (Invitrogen), and DNA was stained with DAPI (Thermo Fisher Scientific). After a final wash in 1× PBS, samples were mounted in FluorSave™ Reagent (Millipore) and covered with a cover glass (Matsunami). Fluorescent images were acquired using a Zeiss LSM900 confocal microscope, with ZEN 3.1 software employed to optimize imaging settings, collect z-stacks, and generate maximum intensity projection images.

### Transmission electron microscopy (TEM) analysis

SGs of wandering L3 larvae were dissected in 1× PBS and fixed overnight at 4 °C in a solution containing 2% PFA and 2% glutaraldehyde in 0.1 M phosphate buffer (pH 7.4). After fixation, samples were washed thrice and post-fixed with 2% osmium tetroxide at 4 °C for 2 hours. The tissues were dehydrated in graded ethanol solutions (50, 50, 90, and 100 %) and were infiltrated with propylene oxide. Samples were then gradually transferred to a fresh 100% resin (Quetol-812; Nisshin EM Co., Tokyo, Japan) and polymerized at 60 °C for 48 hours. Grids were observed under a TEM (JEM-1400Plus; JEOL Ltd., Tokyo, Japan) at an acceleration voltage of 100 kV. Digital images (3296 × 2472 pixels) were captured using a CCD camera (EM-14830RUBY2; JOEL Ltd., Tokyo, Japan). Sample preparation and imaging were performed by Tokai Electron Microscopy, Inc. (Nagoya, Japan).

### Live cell imaging analysis

To visualize microtubules in living cells, larvae expressing β-Tubulin::EGFP were dissected in Schneider’s *Drosophila* medium (SDM). Dissected tissues were incubated in SDM containing 5 mg/mL Hoechst (Sigma) for 30 minutes under dark conditions. After staining, tissues were gently rinsed once with SDM and transferred to the center of a 35 mm glass-bottom dish (IWAKI) along with a drop of SDM (∼20 μl). To prevent floating, insect pins were placed on two sides of the tissues and fixed in position using silicone vacuum grease (Beckman, Brea, CA, USA). Finally, 200 μl of SDM was added on top of the tissues. Images were acquired using a Zeiss LSM900 confocal microscope with a 63× oil immersion objective using the Airyscan mode.

To visualize polytene chromosomes in SGs without fixation, wandering L3 larvae were dissected in SDM and incubated with Hoechst-containing SDM for 30 minutes in the dark. After staining, SGs were mounted in ProLong™ Glass Antifade Mountant and covered with Carl Zeiss™ high-performance cover glasses. Chromosome arms were spread by applying light pressure to the cover glass. Images were acquired with a 40× water immersion objective using the Airyscan mode.

### O-propargyl puromycin (OPP) labeling

OPP labeling was performed following the manufacturer’s guidelines for the Click-iT Plus OPP Alexa Fluor 488 Protein Synthesis Assay kit (Invitrogen) with minor modifications. SGs of wandering L3 larvae were dissected in SDM. A control group was treated with 0.75 mM cycloheximide (Nacalai Tesque, Inc., Kyoto, Japan), a translational elongation inhibitor, in SDM for 2 hours to serve as a negative control, while other groups were harvested in SDM without treatment. Tissues were then transferred immediately to SDM containing 20 μM OPP for 30 minutes. Mock-treated samples, incubated without OPP, served as an additional negative control. Following incubation, tissues were rinsed once with 1× PBS and fixed in 3.7% formaldehyde for 20 minutes. Subsequent steps were carried out according to the kit protocols.

### RNA fluorescence *in situ* hybridization (RNA FISH)

RNA FISH was performed as described previously (Chen et al., 2022; Young et al., 2020). Tissues were dissected from larvae at 72 hAH in 1× PBS and fixed in 3.7% formaldehyde for 30 minutes. After fixation, tissues were washed thrice in 2× SSCT (0.1% Triton X-100 in 2× SSC) for 5 minutes each, then permeabilized in ice-cold 70% ethanol at 4 °C for 30 minutes. Following a brief rinse in 2× SSC with 10% formamide at room temperature, tissues were pre-hybridized for 2 hours at 37 °C in 40 μL hybridization buffer (25% formamide, 1× Denhardt’s solution, 100 mg/mL dextran sulfate, 2× SSC, and 500 μg/mL tRNA). Hybridization was carried out overnight at 37 °C in a thermal cycler (Gene Atlas, ASTEC) with 300 nM probe concentration. Post-hybridization, tissues were washed stepwise for 10 minutes each in prewarmed (37 °C) 40% and then 20% formamide in 2× SSCT, followed by three 10-minute washes in 2× SSCT and a 10-minute wash in 0.1× PBT (0.1% Triton X-100 in 1× PBS). Samples were counterstained with DAPI for 20 minutes, then washed twice in PBS. Samples were then mounted in ProLong™ Glass Antifade Mountant, and coverslipped with Carl Zeiss™ high-performance cover glasses. Imaging was performed immediately using a Zeiss LSM900 confocal microscope with a 40× water immersion objective.

### Re-evaluation of chromosomal structure in prothoracic gland cells

Histochemical images of PG cells from 159 RNAi animals, previously characterized by enriched DNA staining, interspaces in nuclei, or multinucleolar-like structures (classified as groups A, C, and D, respectively; Ohhara et al., 2019), were manually re-evaluated using visual, qualitative criteria. PG cells displaying a distinct DNA-enriched “blob”-like chromosomal structure were classified as positive hits. Cells showing moderate changes in chromosomal morphology without a clear “blob”-like structure were categorized as having a “subtle alteration of chromosomal structure”. Cells lacking any condensed DNA staining pattern were designated as having “no obvious phenotype”.

### LysoTracker staining

Acidic autolysosomes were visualized using LysoTracker as described (Martelli, 2023; Mauvezin et al., 2014). SGs from wandering L3 larvae were dissected in SDM and incubated with LysoTracker™ Red DND-99 (Invitrogen) in the dark for 30 minutes at room temperature. Following incubation, samples were washed thrice with 1× PBS for 5 minutes each and then fixed in 3.7% formaldehyde for 30 minutes. After fixation, tissues were washed thrice with 1× PBS for 5 minutes each. DNA was counterstained with DAPI.

### Western blotting

Eight to nine wandering L3 larvae were homogenized in 1× PBS. Lysates were mixed with an equal volume of 2× SDS sample buffer (4% SDS, 20% Glycerol, 0.004% bromophenol blue, and 0.125 M Tris-HCl) containing 10% 2-mercaptoethanol (nuclease tested) and heated at 95°C for 5 minutes to denature. Samples were separated by 15% PAGE and transferred to a PVDF membrane. The membrane was blocked with 0.3% skim milk in TBST (1× Tris-buffered saline with 0.1% Tween-20) at room temperature for 1 hour. Membranes were incubated overnight at 4 °C with primary antibodies—mouse anti-FLAG (Sigma-Aldrich) and mouse β-Actin Antibody (Santa Cruz Biotechnology)—in 0.3% skim milk containing 10% immunoreaction enhancer solution 1 (TOYOBO). After washing thrice with TBST, membranes were incubated for 2 hours at room temperature with HRP-conjugated anti-mouse IgG secondary antibody (Cytiva) in 0.3% skim milk containing 10% immunoreaction enhancer solution 2 (TOYOBO). Protein bands were detected using SuperSignal™ West Atto Ultimate Sensitivity Substrate (Thermo Scientific) and imaged with a chemiluminescence detector CCD camera (ImageQuant 800).

### Northern blotting

RNA was extracted from SGs of wandering L3 larvae using RNAiso Plus. Equal amounts of RNA (2 µg per lane) were separated on a 1.2% formaldehyde-agarose gel (1.2% agarose, 1× MOPS buffer, 2% formaldehyde) alongside the ^DynaMarker^RNA High Ladder (BioDynamics Laboratory, Pittsburgh, PA) as a size marker. After electrophoresis, marker bands were visualized with 1 mg/mL ethidium bromide under UV light. RNA was then transferred overnight onto a nitrocellulose membrane (Macherey-Nagel, Düren, Germany) and crosslinked by 120 mJ/cm² UV irradiation using DNA-FIX (ATTO). Membranes were pre-hybridized for 30 minutes at 68 °C in DIG Easy Hyb solution (DIG Northern Starter Kit) with gentle agitation, followed by overnight hybridization at 68 °C with 1 µg of DIG-labeled 28S rRNA antisense probe. Post-hybridization washes included two 5-minute rinses in prewarmed stringency wash buffer I (2× SSC, 0.1% SDS) at room temperature, two 15-minute washes in prewarmed stringency wash buffer II (0.1× SSC, 0.1% SDS) at 68°C, and equilibration in washing buffer (0.3% Tween 20 in maleic acid buffer) for 2 minutes at room temperature. Membranes were blocked with 1× blocking solution (DIG Northern Starter Kit) in maleic acid buffer (0.1 M maleic acid, 0.15 M NaCl, pH 7.5) for 30 minutes, then incubated with DIG antibody (DIG Northern Starter Kit) diluted in 1× blocking solution for 30 minutes at room temperature, followed by two 15-minute washes in washing buffer. For signal detection, membranes were incubated in detection buffer (0.1 M Tris, 0.1 M NaCl, pH 9.5) for 5 minutes before adding the chemiluminescent substrate (DIG Northern Starter Kit) for an additional 5 minutes. Membranes were covered with UV-transparent plastic wrap and exposed for 10 minutes using a chemiluminescence detector CCD camera (ImageQuant 800).

### RNA-seq

For each replicate, 10 pairs of SGs in wandering L3 larvae were collected from control samples, and 30 pairs were collected from each of the four RNAi-treated groups. Quality control, read mapping, and counting were performed by Tsukuba i-Laboratory LLP. Differential gene expression and GO enrichment analysis were carried out in the R software environment. Differentially expressed genes were determined using the *DESeq2* R package, with criteria set at log_2_ fold-change > 1 and adjusted *p*-value < 0.05. The R package *clusterProfiler* was used to identify enriched GO terms. All plots were constructed using the *ggplot2* R package.

### Statistical measurements

Fluorescence intensity and area were measured using Fiji/ImageJ (https://fiji.sc). The size of SG was quantified by measuring the F-actin-stained area of each gland lobe following Phalloidin-Alexa-Fluor labeling. Nuclear size was assessed by outlining the nuclear envelope marker Lamin C. NudC signals in the nucleolus were quantified by extracting NudC::GFP-positive signals co-localized with the nucleolar marker Fib. γH2Av signal quantification involved extracting γH2Av-positive nuclear signals that overlapped with DAPI.

Statistical analysis and data visualization were performed using GraphPad Prism 10.5.0. Differences between two groups were assessed by the Mann–Whitney test. Comparisons among three or more groups were evaluated using Brown-Forsythe and Welch ANOVA tests, followed by Dunnett’s T3 multiple comparisons test. Sample sizes, randomization, and blinding were not applied, as qualitative phenotypes were markedly distinct from wild type. Results are presented as mean values with standard deviation. Statistical significance was defined as *p* < 0.05*, < 0.01**, < 0.001***, and < 0.0001****. Figures were assembled in Adobe Illustrator 2025 (v29.0).

### Raw data availability

All numerical raw data used in the main and supplemental figures are described in the Supplemental Source Data file. All of the raw RNA-seq data reported in this study were deposited in the Sequence Read Archive at DNA Data Bank of Japan (DRA; BioProject PRJDB35869 with Run DRR717927-DRR717932 for comparison between control and *NudC* RNAi; and BioProject PRJDB35873 with Run DRR717933-DRR717944 for comparison of control, *eIF5* RNAi, *RpLP0-like* RNAi, and *Nopp140* RNAi).

## Supporting information

Supplemental Figures S1 to S11, and their figure legends

Legends for Supplemental Tables S1, S2, and S3

Supplemental Source Data for all numerical data represented in figures

## Acknowledgements

We thank Fumiaki Obata, Michael B. O’Connor, Naoki Yamanaka, Developmental Studies Hybridoma Bank, Bloomington Drosophila Stock Center, Kyoto Stock Center, Tsinghua Fly Center, and Vienna Drosophila Resource Center for providing materials and reagents used in the experiments. We are also grateful to Shintaro Iwasaki, Minoru Moriyama, and Yuichi Shichino for technical assistance; Daiki Kitamura for critical reading of the manuscript; and Editage (www.editage.com) for editing and reviewing this manuscript for English language. This work was supported by the JST SPRING Grant Number JPMJSP2124 to D.S., the Japan Society for the Promotion of Science (JSPS) (KAKENHI) Grant Number 24K09466 and 21H02521 to Y.S.-N. and Y.O., respectively, MEXT Supported Program for the Inter-University Research Network for High Depth Omics, Institute of Molecular Embryology and Genetics (IMEG), Kumamoto University to A.N., National Natural Science Foundation of China (32070499) to W.S., and the program of the Joint Usage/Research Center for Developmental Medicine, IMEG, Kumamoto University (K23-06 and K24-15) to R.N.

