## Supplemental Figures S1 to S11, and their figure legends for "NudC moonlights in ribosome biogenesis and homeostasis in *Drosophila melanogaster* polyploid cells"

### Supplemental Figure Legends

#### Figure S1

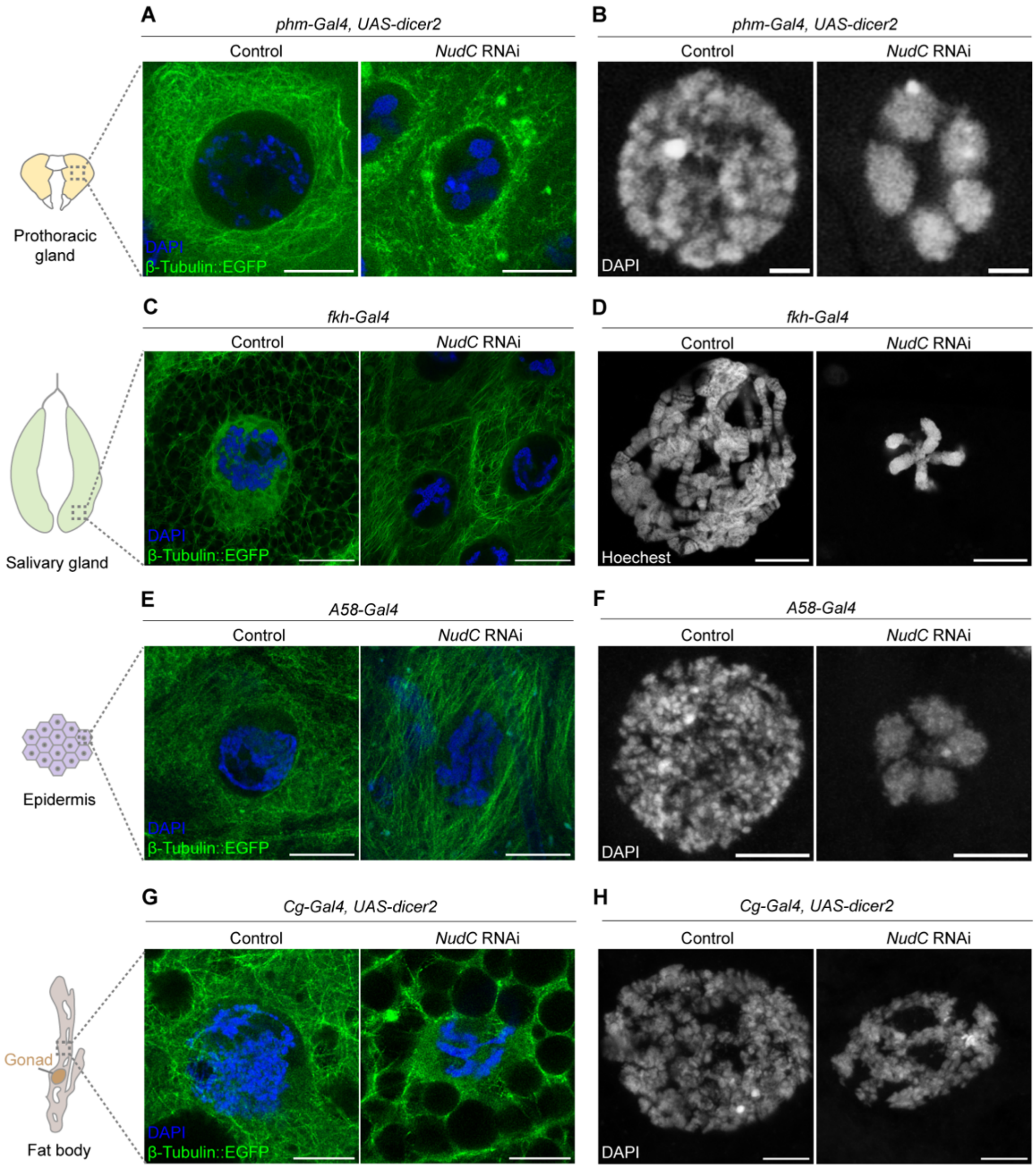


**Figure S1. *NudC* knockdown disrupts microtubule organization and chromosome structure in polyploid cells.**
Analysis of control and *NudC* RNAi polyploid cells, including the prothoracic gland (A and B), salivary gland (C and D), epidermis (E and F), and fat body (G and H). Figures (B) and (D) correspond to Figure 1E and Figure 2I, respectively.

(A, C, E, G) Microtubules were visualized using β-Tubulin::EGFP (green) with DNA counterstained by DAPI (blue). Microtubule organization was particularly difficult to assess in the fat body due to the abundance of storage lipids.
(B, D, F, H) Polytene chromosomes were stained with DAPI (white); in SG cells, Hoechst staining was used, and chromosome arms were gently spread.
Scale bars: 10 μm (A), 2 μm (B), 20 μm (C), 10 μm (D), 10 μm (E), 5 μm (F), 10 μm (G), 5 μm (H).

### Figure S2

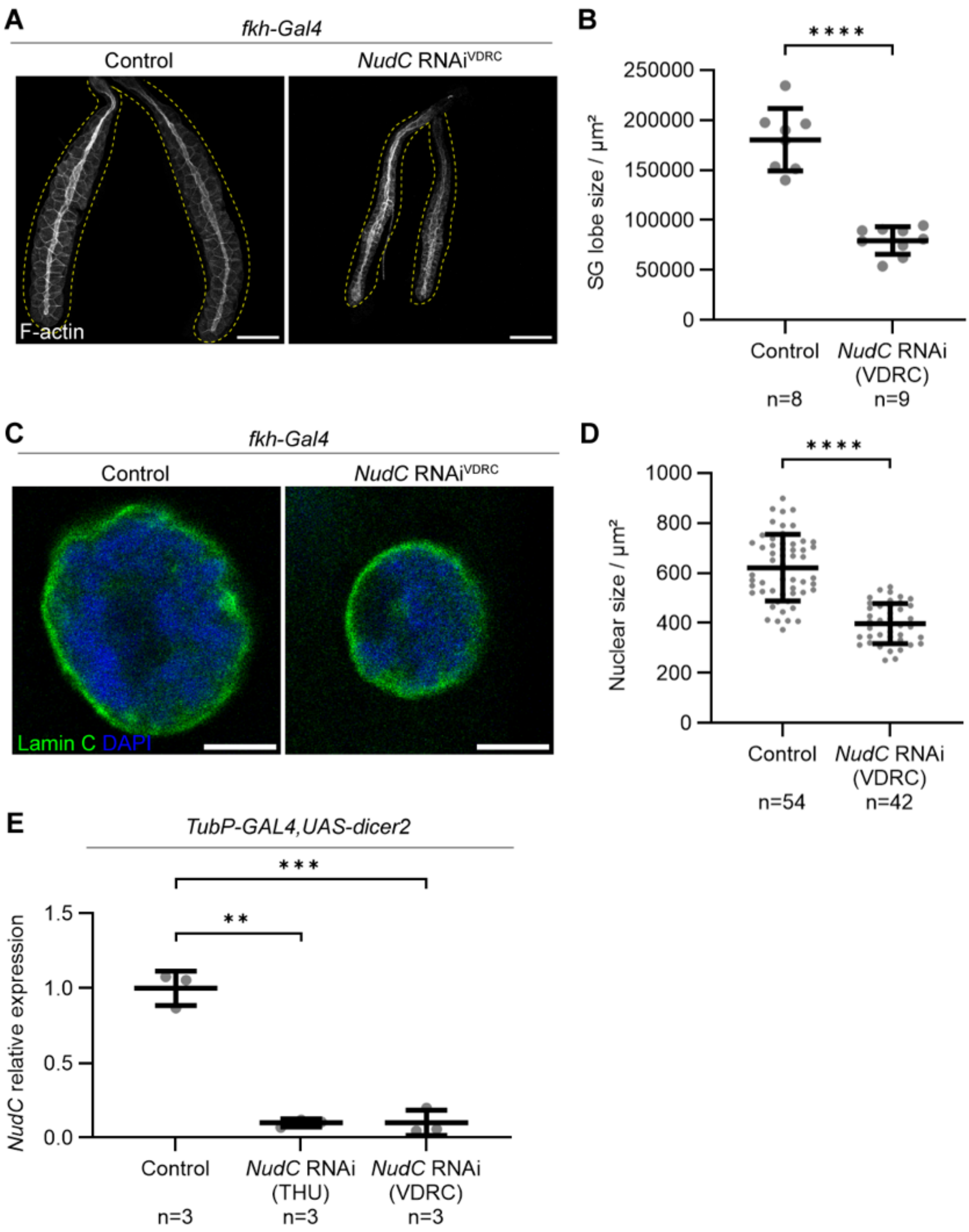


**Figure S2. *NudC* knockdown using a second RNAi strain recapitulates SG defects.**
(A and B) SG lobe size in control (*fkh*>+) and *NudC* RNAi$^{VDRC}$ (*fkh>NudC RNAi$^{VDRC}$*) larvae at the wandering L3 stage. (A) Representative SGs stained with F-actin (white) and outlined with yellow dashed lines. Scale bar: 200 µm.

(B) Quantification of SG lobe area from control (n = 8) and *NudC* RNAi (n = 9) larvae. Bars show mean ± SD. ****$p < 0.0001$, Mann–Whitney test.
(C and D) Nuclear size in control and *NudC* RNAi$^{VDRC}$ SG cells. (C) Representative images. Green, Lamin C; Blue, DAPI. Scale bar: 10 µm. (D) Nuclear area measured in control (n = 54) and *NudC* RNAi$^{VDRC}$ (n = 42) SG cells. Six cells per SG lobe were randomly selected for measurement in each genotype. Bars show mean ± SD. ****$p < 0.0001$, Mann–Whitney test.
(E) Relative *NudC* expression levels in two independent *NudC* RNAi strains (*tubP>dicer2, NudC RNAi$^{THU}$* and *tubP>dicer2, NudC RNAi$^{VDRC}$*) compared with control (*tubP>dicer2,* +). Bars represent mean ± SD. **$p < 0.01$, ***$p < 0.001$ (Brown–Forsythe and Welch ANOVA, followed by Dunnett's T3 multiple comparisons)

### Figure S3

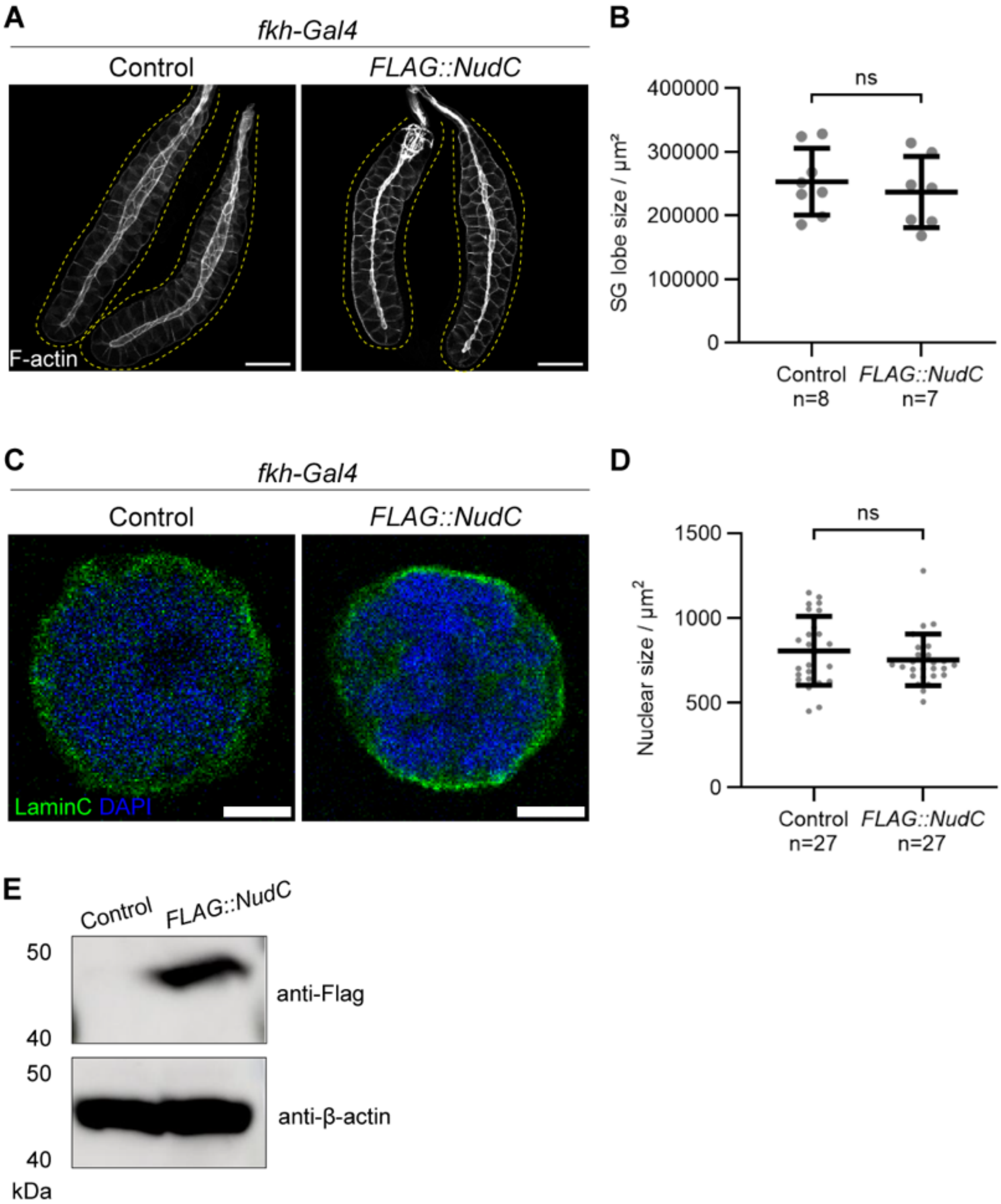


**Figure S3. Overexpression of *NudC* does not promote the SG tissue or nuclear growth.**

(A and B) SGs from control and *FLAG::NudC* larvae at the wandering L3 stage. (A) Representative SGs stained with F-actin (white) and outlined with yellow dashed lines. Scale bar: 200 μm. (B) Quantification of SG lobe area in control (n

= 8) and *FLAG::NudC* (n = 7) larvae. Bars show mean ± SD. No significant difference (ns), Mann–Whitney test.
(C and D) Nuclear size in control and *FLAG::NudC* SG cells. (C) Representative images showing Lamin C (green) and DAPI (blue). Scale bar: 10 μm. (D) Quantification of nuclear area from control (n=27) and *FLAG::NudC* (n = 27) larvae. Bars represent mean ± SD. No significant difference (ns), Mann–Whitney test.
(E) Detection of FLAG-tagged NudC by western blotting. Lysates were prepared from wandering L3 larvae expressing FLAG::NudC under the tubP-GAL4 driver (*tubP>dicer2, FLAG::NudC*). Control samples lacked the transgene (*tubP>dicer2,* +). β-Actin served as a loading control.

**Figure S4**

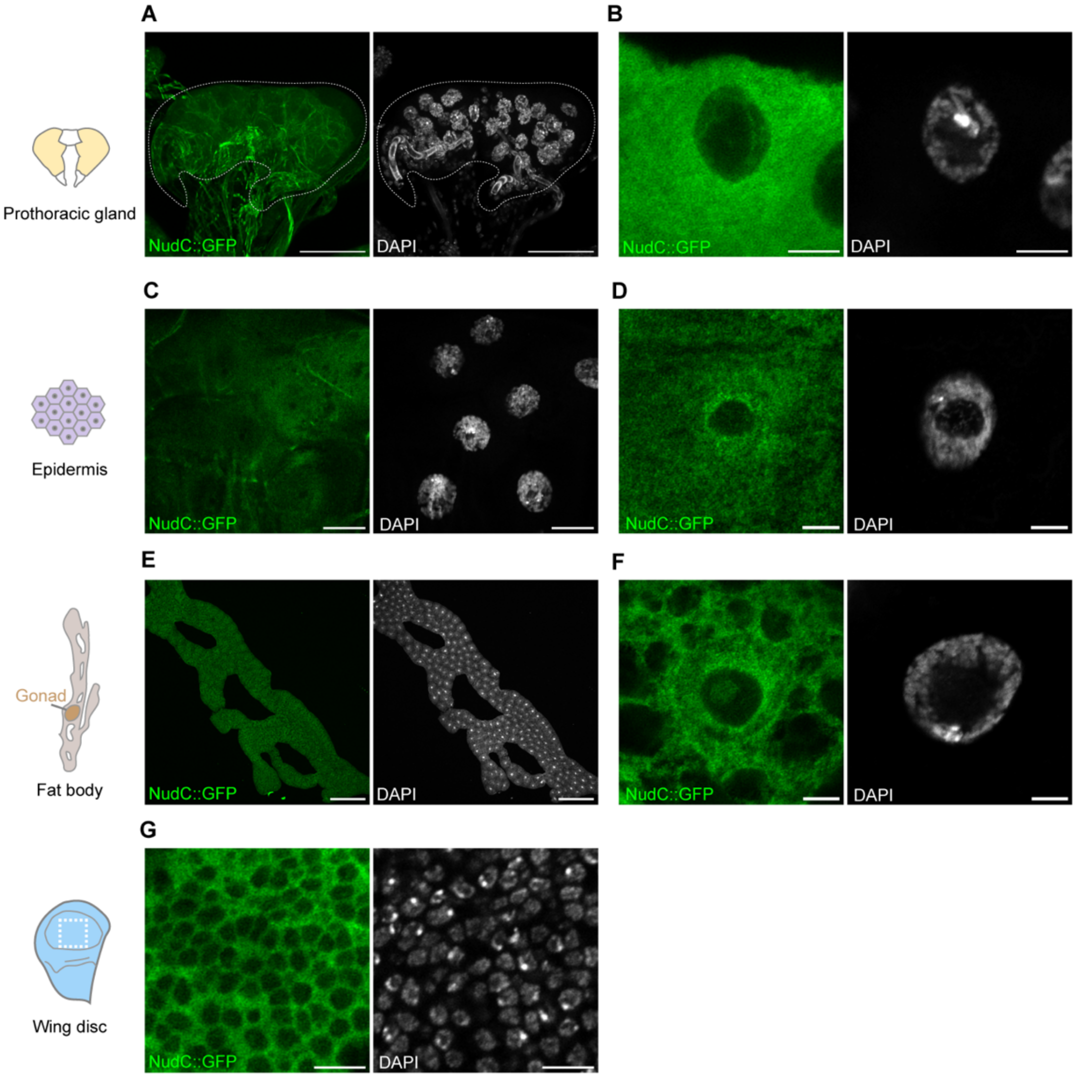


**Figure S4. Expression and subcellular distribution of NudC::GFP in multiple larval tissues.**

Localization of NudC::GFP in the prothoracic gland (A and B), epidermis (C and D), fat body (E and F), and imaginal wing disc (G) from the wandering L3 larvae. Low- (A, C, E) and high-magnification (B, D, F) views are shown. In (E and F), fat body cells adjacent to the gonads are displayed. Scale bars: 50 μm (A), 5 μm (B), 20 μm (C), 5 μm (D), 200 μm (E), 5 μm (F).

Figure S5

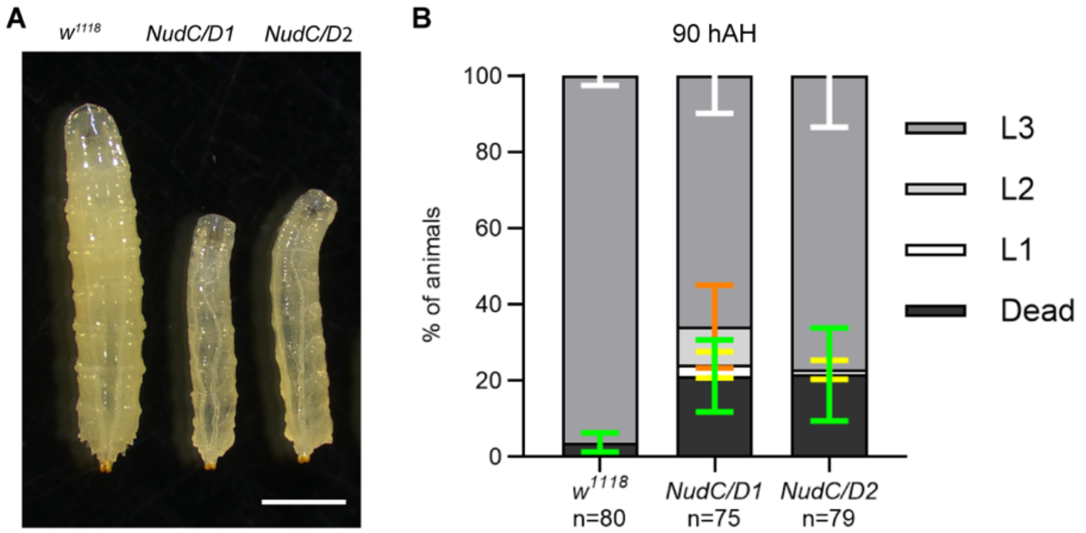


**Figure S5. Survival rate and developmental stages of wild-type and *NudC* transheterozygous mutants.**

(A) Larval body size of wild-type (*w*$^{1118}$) and two *NudC* transheterozygous mutants (*NudC*$^{GS15156}$/ *NudC*$^{Df(3L)ED223}$, *NudC/D1*; *NudC*$^{GS15156}$/ *NudC*$^{Df(3L)ED4674}$, *NudC/D2*) at 90hAH. Scale bar: 1 mm.

(B) Survival and developmental progression of *w*$^{1118}$ and *NudC* mutant larvae (n = 75–80). Larval instar stages were determined by anterior spiracle morphology (Koyama and Mirth, 2021). Each genotype was divided into four groups, shown as dead larvae (green), L1 (yellow), L2 (orange), and L3 (white) error bars.

### Figure S6

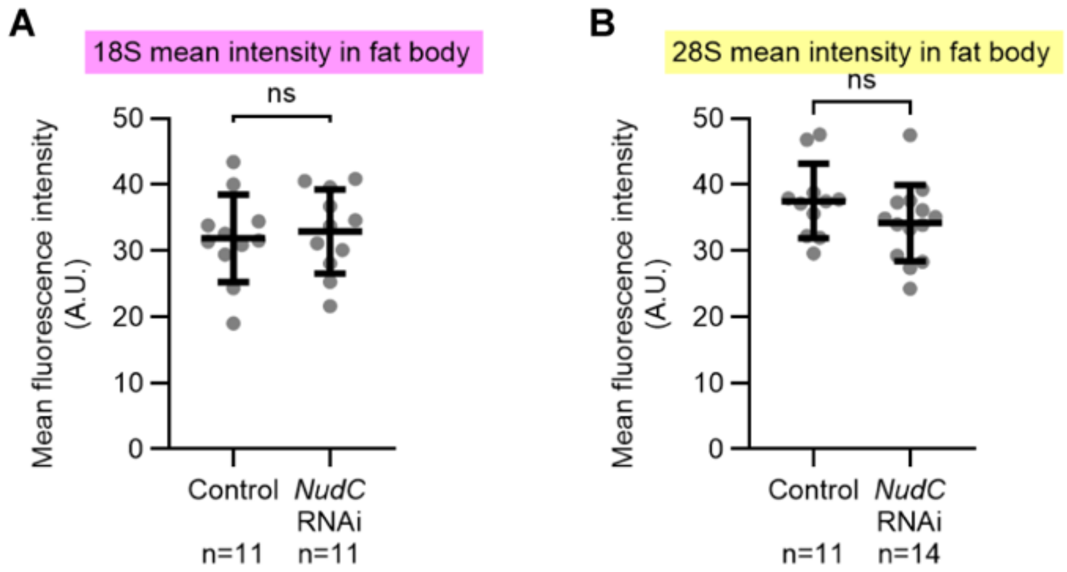


**Figure S6. rRNA signals in fat body cells surrounding control and *NudC* RNAi SGs.**
Scatter plots of mean fluorescence intensity for 18S (A) and 28S (B) rRNA FISH probes in fat body cells adjacent to control or *NudC* RNAi SGs (n = 11–14). Bars show mean ± SD. No significant difference (ns), Mann–Whitney test.

Figure S7

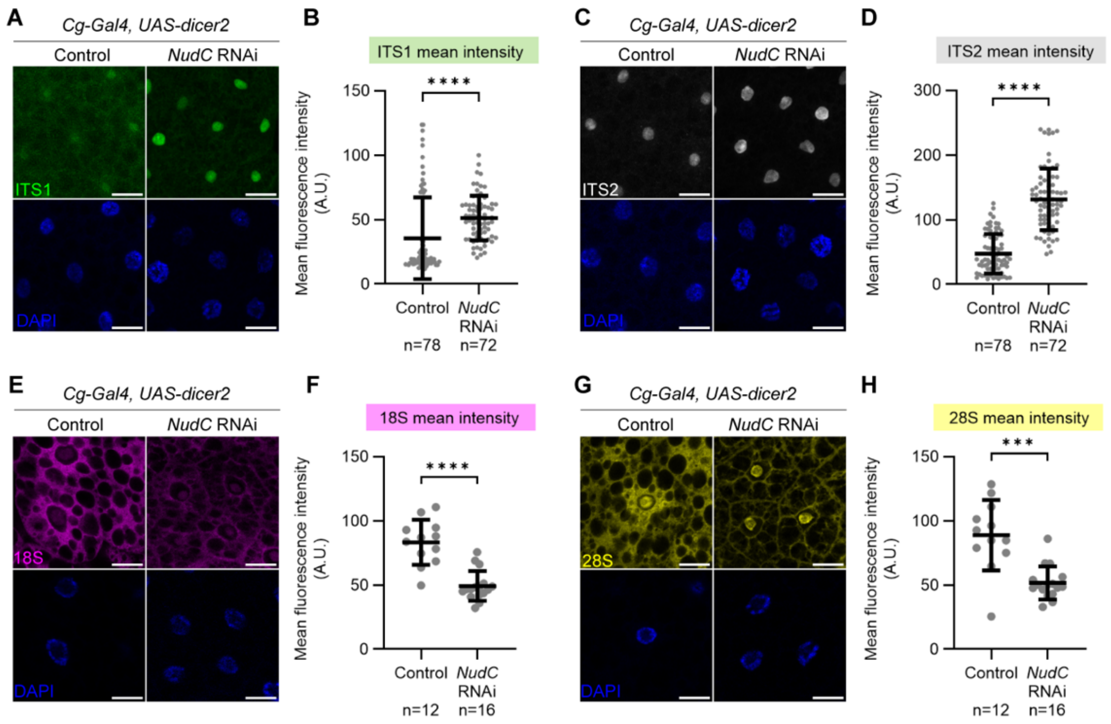


**Figure S7. *NudC* is required to maintain rRNA levels in the fat body.**
RNA FISH in fat body cells of L3 male larvae at 72 hAH. Probes targeted ITS1 (A and B; green), ITS2 (C and D; white), 18S (E and F; magenta), and 28S (G and H; yellow). DNA was counterstained with DAPI (blue). Scale bar: 20µm. (B, D, F, and H) Quantification of FISH signal intensities. ITS1 (B) and ITS2 (D) signals were measured in individual cells (n = 72–78). 18S (F) and 28S (H) signals represent average fluorescence per defined region adjacent to the gonads from each fat body sample (n = 12–16). Bars show mean ± SD. Mann–Whitney test, ***$p$ < 0.001, ****$p$ < 0.0001.

Figure S8

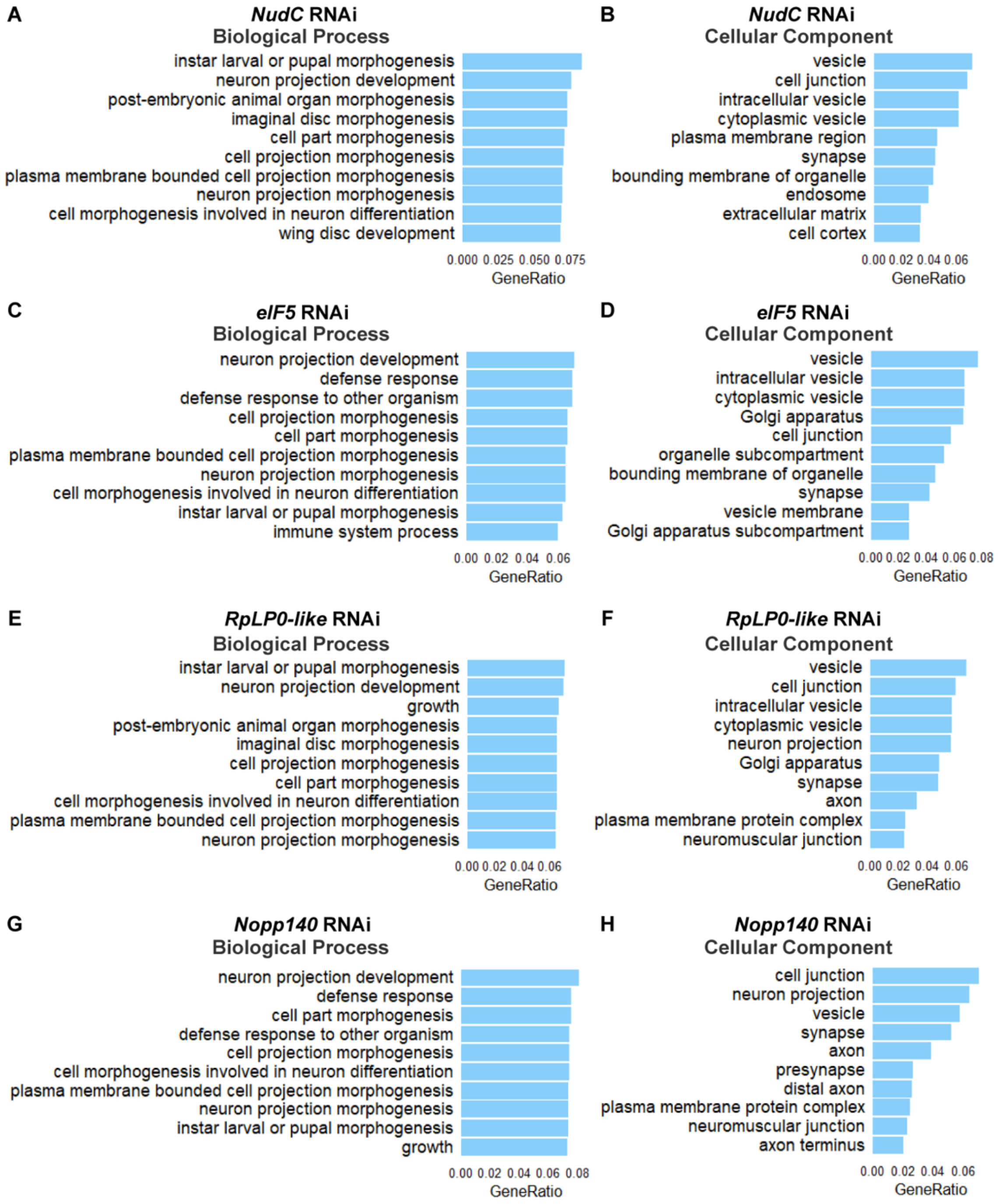


**Figure S8. GO enrichment of down-regulated DEGs upon knockdown of *NudC* or RBFs in SGs.**

Top 10 GO terms ($p < 0.05$) for down-regulated genes in SGs of wandering L3 larvae upon knockdown of *NudC* (A and B), *eIF5* (C and D), *RpLP0-like* (E and F), or *Nopp140* (G and H). Terms are shown for biological process and cellular component categories. Bars indicate the ratio of down-regulated genes per category.

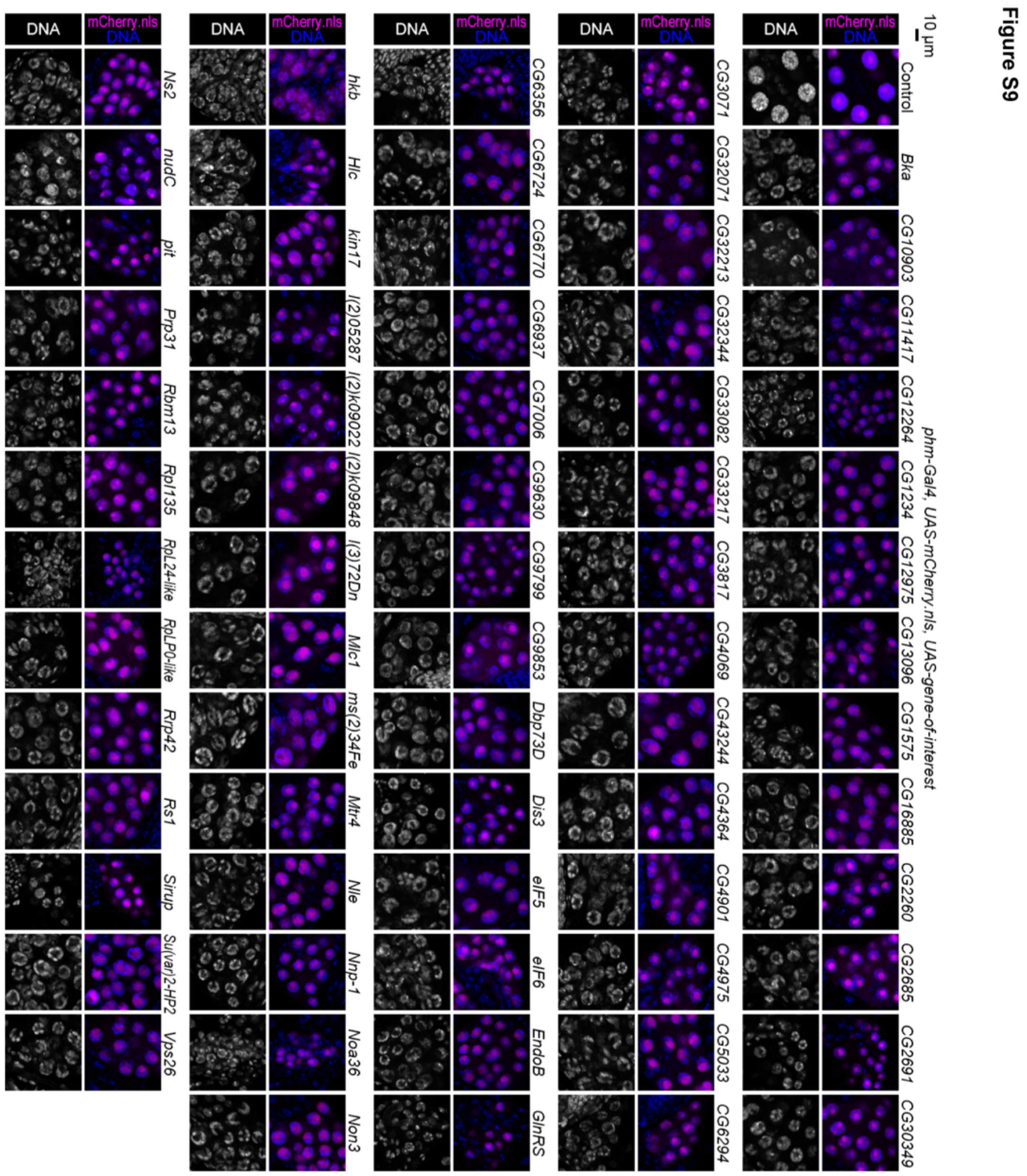


**Figure S9. DNA-enriched "blob" chromosomal structure identified in RNAi screening of PG cells.**

Representative PG cells from control (*phm>mCherry.nls*) and RNAi larvae (*phm>mCherry.nls, RNAi against gene-of-interest*) showing the "blob" chromosome structure. DNA was stained by Hoechst (blue and white in the upper and lower panels, respectively), and the nuclei of PG cells were labelled by mCherry.nls (magenta in the upper panels). Images are from the PG-selective RNAi screen (Ohhara et al., 2019) and re-evaluated as the positive hits with the "blob" chromosome structure.

**Figure S10**

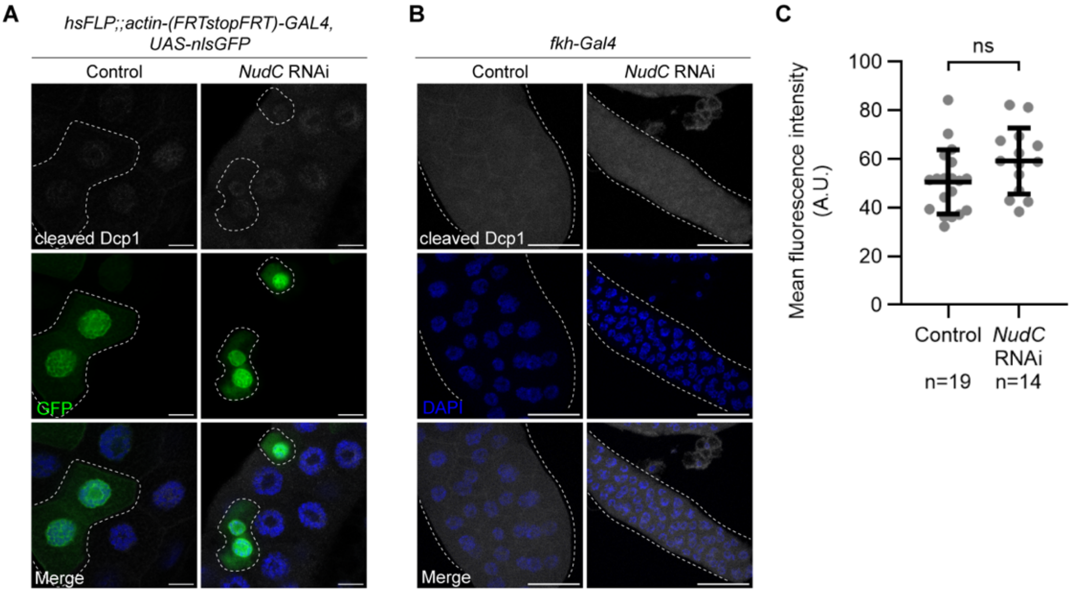


**Figure S10. *NudC* knockdown does not significantly increase apoptosis in SGs.**

(A) Mosaic SG clones from control or *NudC* RNAi larvae stained with anti-cleaved Dcp1 (white). Clones are marked by GFP and dashed outlines. DNA was counterstained with DAPI (blue). Scale bar: 20 μm.

(B) Whole SGs immunostained with anti-cleaved Dcp1 (white) in control and *NudC* RNAi wandering L3 larvae. Scale bar: 100 μm.

wandering larvae. Scale bar: 100 μm.

(C) Quantification of anti-cleaved Dcp1 fluorescence intensity in SGs (control, n = 19; *NudC* RNAi, n = 14). Bars show mean ± SD. Mann–Whitney test: not significant (ns > 0.05).

**Figure S11**

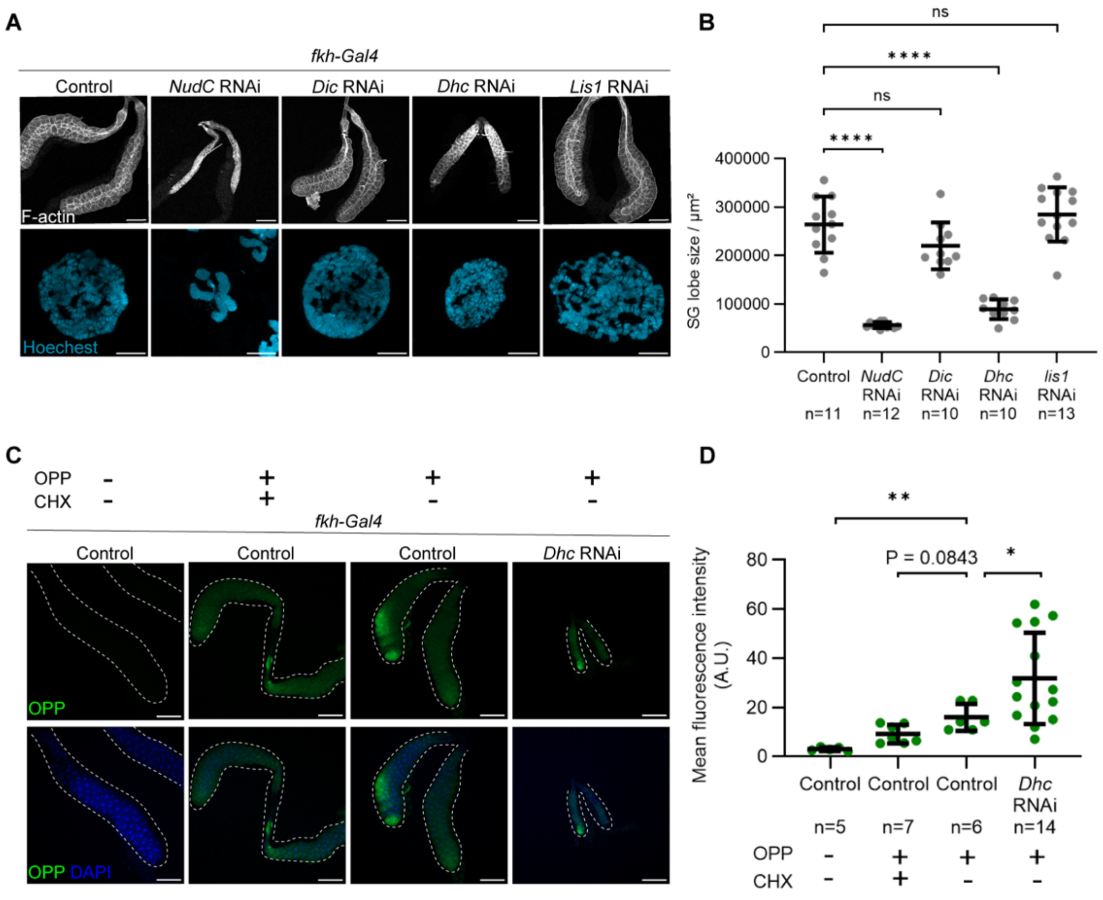


**Figure S11. *NudC* RNAi phenotype is distinct from knockdown of dynein-related genes.**

(A) SGs (upper panels) and polytene chromosomes from SG nuclei (lower panels) of control, *NudC* RNAi, *Dynein intermediate chain* (*Dic)* RNAi, *Dynein heavy chain* (*Dhc*) RNAi, and *Lis1* RNAi at the wandering L3 stage. Scale bars: 200 µm (upper), 10 µm (lower).

(B) Quantification of SG lobe size. Sample sizes: control (n = 11); *NudC* RNAi (n = 12); *Dic* RNAi (n = 10); *Dhc* RNAi (n = 10); *Lis1* RNAi (n = 13). Bars show mean ± SD. ****$p < 0.0001$; ns > 0.05 (Brown–Forsythe and Welch ANOVA followed by Dunnett's T3 multiple comparisons test).

(C and D) Protein synthesis assayed by OPP signals in control and *Dhc* RNAi SGs at the wandering L3 stage. SGs (outlined by dashed lines) were incubated with or without OPP; CHX pretreatment served as a negative control. (C) Representative images of OPP signals (green); DNA counterstained with NuclearMask™ Blue. Scale bar: 200 µm. (D) Quantification of OPP fluorescence intensity (n = 5, 7, 6, 14 SGs/group). Bars show mean ± SD.

$^{*}p < 0.05$, $^{**}p < 0.01$; ns > 0.05 (Brown–Forsythe and Welch ANOVA with Dunnett's T3 test).
