## Supplementary material for "NudC moonlights in ribosome biogenesis and homeostasis in *Drosophila melanogaster* polyploid cells": Legends for Supplemental Tables S1, S2, and S3

**Supplemental Table legends**

**Table S1. Re-evaluation of chromosomal structure in PG cells.**

* BDSC, Bloomington Drosophila Stock Center; BDSC TRiP, Transgenic RNAi Project in BDSC; VDRC, Vienna Drosophila Resource Center; NIG, National Institute for Genetics.

** A, enriched DNA staining; C, interspaces in nuclei; D, multinucleolar-like structures.

(XLSX)

**Table S2. GO enrichment analysis of 68 hit genes.**

GO enrichment was performed using the *GO.db* R package. For the four unidentified genes, updated information was retrieved from InterPro (Blum et al., 2025) and FlyBase (release FB2025_02). Gene symbols in bold denote RBFs. Genes selected for this study are highlighted in yellow.

(XLSX)

**Table S3. Insertion sequence used to generate the NudC::GFP knock-in strain.**

(XLSX)
